# Genetic Influences on Educational Attainment Through the Lens of the Evolving Swedish Welfare State: A cross-level gene-environment interaction study based on polygenic indices and longitudinal register data

**DOI:** 10.1101/2023.11.02.565287

**Authors:** Oskar Pettersson

## Abstract

Gene-environment interaction with regards to educational attainment has received increasing attention during the last few years. However, the potential interdependence between different types of environments in gene-environment interaction models has mostly been neglected. Using high-quality register data for an extensive panel of Swedish twins, born during most of the twentieth century, this study explores how genetic propensities for educational attainment, as measured by a polygenic index, interact with both macro-level institutional and sociopolitical context, and with socioeconomic background. The analyses, which combine between-family and causally robust within-family models, suggest that the average association between genetic propensities and educational attainment has increased in Sweden during the twentieth century, along with the expansion of the educational system and decreased economic inequality. There is also evidence of a positive interaction between genetic propensities and socioeconomic background, but only in the oldest cohorts in the sample, and that were born before the Swedish welfare state had been fully established. This implies that micro-level gene-environment interactions can be significantly dependent on macro-level context, an insight that has arguably not yet been given sufficient attention in the literature. Acknowledging limitations of polygenic indices, and the arbitrariness of the genetic lottery, the results may nevertheless indicate a development towards higher equality of opportunity in Sweden during the twentieth century.

## 1 Introduction

It has long since been established that we live in the age of the *knowledge economy*, where the importance of education has increased substantially (Esping-Andersen 2005). Indeed, educational attainment is an increasingly important indicator of life-course achievement and status, and is strongly related to various other outcomes, such as earnings, occupation, and health. On the level of policy making, investing in education is also regarded as a solution to a variety of structural challenges (Morel et al. 2011). Understanding what can explain individual-level educational outcomes, then, is crucial for social scientists and policy makers alike.

Educational attainment, often (but not always) defined as the number of years in formal education, has traditionally been understood through the lens of parental (e.g. Breen and Jonsson 2005) and neighbourhood (e.g. Chetty et al. 2016) socioeconomic status. However, twin and adoption studies have shown that *genetic* factors, too, explain a substantial amount of the variation in educational attainment (Branigan et al. 2013; Silventoinen et al. 2020). Recent genome-based studies have also begun mapping the multiple genetic variants that, albeit in a distal sense, go on to influence educational attainment (e.g. Okbay et al. 2022).

While both genetic factors (here understood as underlying *genetic propensities* for education) and social environments should be expected to be important for educational attainment in their own right – each adding to a fuller explanation – it is crucial to explore the interplay between them. Indeed, the simple dichotomy between ’nature’ and ’nurture’ has long since been deemed obsolete (Turkheimer 2000). One specific type of gene-environment interplay is gene-environment *interaction*: that genes have different effects depending on different environmental characteristics (Plomin et al. 1977). Only quite recently, researchers from across the social sciences have begun applying genomic research methods to investigate this topic, often using *polygenic* indicators of genetic propensity. Researchers have asked, for example, whether genetic influences on educational attainment are moderated by important micro-level environments like socioeconomic background (e.g. Papageorge and Thom 2020), meso-level environments like schools and neighbourhoods (e.g. Cheesman et al. 2022), and, but to a far lesser extent, structural or macro-level environments like countries or time-periods (e.g. Lin 2020).

Apart from having mostly overlooked the potentially important moderating role of macro-level environments, this literature has often investigated specific types of environments in isolation, thereby leaving the potential *interdependencies* between different scales of environments unexplored (Boardman et al. 2013). The aim of this paper is to help address these two shortcomings. First, it investigates whether the average magnitude of genetic influences on educational attainment, as captured by a polygenic index for educational attainment (an individual-level genetic predictor that captures genetic variants associated with educational attainment) differs depending on differences in *temporal* macro-level context. Second, it investigates whether the same genetic influences are moderated by another important type of environment, namely socioeconomic background, but with the key assumption that this micro-level gene-environment interaction may itself be moderated by macro-level context. I will argue that studying both these dimensions in tandem is not only theoretically sound, but also crucial from the perspective of equality of opportunity.

The paper studies Sweden during the twentieth century. In this period, Sweden underwent significant economic, social and educational developments, going from being one of the poorer countries in Europe to one of its wealthiest, being now consistently found to excel in terms of equality of opportunity and social mobility (Breen and Jonsson 2007). Specifically, the study analyses a noteworthy sample of genotyped twin cohorts in the Swedish Twin Registry (STR), born between the 1920’s and the 1990’s, and that can be linked to high-quality register data on educational attainment, as well as other related outcomes. This sample provides a hitherto unprecedented opportunity to examine genetic influences on educational attainment through the lens of macro-level context, with birth year differences so large as to capture potentially quite substantial macro-level differences within the same country. Indeed, previous research has mainly been able to study cohort moderation in the earlier decades of the twentieth century. Notably, the registers also enables parental education to be measured for cohorts born as early as 1940. This allows for the particular contribution of showing whether a potential interaction between genetic factors and socioeconomic background is itself dependent on macro-level developments. Finally, the twin sample also contains a sizeable number of dizygotic twin pairs, which allows for causally robust within-family analyses, the call for which is regularly being made in the emerging social science genomics literature (Cheesman et al. 2023; Harden and Koellinger 2020).

The results of the paper suggest the following: (1) the association between genetic propensities for educational attainment and actual educational attainment has increased in Sweden during the twentieth century; (2) said association is stronger for twins with more highly educated parents, but only among the oldest twins in the sample containing parental information. Indeed, the *complementary* interaction between genetic propensities and socioeconomic background that exists for the oldest cohorts appears to have decreased over time. Not only, then, do genetic influences on education appear to increase as the macro-level environment becomes more favourable, but the socioeconomic stratification of the effects also appears to decrease. These results also extend to a set of supplementary outcomes. Acknowledging limitations related to the use of polygenic indices, as well as the arbitrariness of the genetic lottery, the results may nevertheless indicate a development towards higher equality of opportunity in Sweden during the twentieth century.

## 2 Theory and previous research

Gene-environment interaction, then, refers to a phenomenon in which the effects of genetic propensities on an outcome differ depending on the environment to which said genetic propensities are exposed (Plomin et al. 1977). This begs an important question of how the environment should be understood and defined. This paper takes as its theoretical starting point a type of definition appropriately formulated by Boardman et al. (2013), in which the environment is an ”external, multilevel, and multidimensional feature that determines an individual’s exposure to risks and access to resources” (p. 65). Being multi-levelled, the environment thus operates on the micro-level (families), meso-level (schools, neighbourhoods, etc.), and even on the macro-level (countries and time-periods). Being multidimensional, moreover, it encompasses physical, social, economic, and institutional dimensions – all of which contribute to making certain environments more abundant in resources – and thereby more *favourable* – which in turn can moderate the role played by genes (Boardman et al. 2013). For the purposes of this study, the environment is simplified as having a macro and a micro level. In the paragraphs below, I further define the micro- and macro-level environment from the perspective of resources, and derive expectations regarding their moderating effects on genetic influences on educational attainment. To this is then added the possibility that these environmental levels can interact with each other, in a more complex model of gene-environment interaction.

While there are many different dimensions that could be mentioned, a favourable *macro*-level environment can in this context be appropriately understood as a structural, sociopolitical and institutional context characterised first and foremost by high economic development, high educational spending, and low economic inequality (Adkins and Vaisey 2009; Baier et al. 2022; Selita and Kovas 2019). The institutional design of the educational system can also have important equalizing effects, with one important dimension being the degree to which the system postpones the sorting of students into vocational and academic tracks (Brunello and Checchi 2007; Pfeffer 2008). Adequate access to pre-schooling can also be mentioned in this context, having indeed been described as “possibly far more decisive than education reforms in terms of equalizing the opportunity structure” (Esping-Andersen 2015, p. 126).

A favourable *micro*-level environment can straightforwardly be defined in terms of socioeconomic background, or, indeed, parental socioeconomic status. To have adequate access to economic, social and cultural resources within the family environment may be crucial for an individual’s educational career. Parents with high socioeconomic status can provide not only concrete, material resources – money, adequate living conditions, a safe neighbourhood, access to good schools, and so on – but also social and cultural investments that are often deemed crucial for children’s educational careers, such as motivation and involvement in the child’s education (Breen and Jonsson 2005; De Graaf et al. 2000; Erola et al. 2022).

By shaping individuals’ access to various resources, both the macro- and micro-level environment are here expected to be important moderators of the effects of education-favourable genetic propensities on actual education. The next key question is how these environments should be expected to moderate these genetic influences. One quite influential idea from the developmental behavioural genetics literature is that a favourable environment would serve to *increase* genetic influences. As argued by Shanahan and Hofer (2005), they can ’enhance’, or, in the terminology adopted here, *complement* genetic propensities. Although Shanahan and Hofer discuss this complementary mechanism in a micro- or possibly meso-level context, a favourable macro-level environment should also have the potential to have direct, complementary moderating effect on genetic influences; if economic development and education spending is high, and if inequality is low, genetic factors could be allowed to express themselves in a way that they would not in a less favourable macro-level context (Adkins and Vaisey 2009; Selita and Kovas 2019). For example, Branigan et al. (2013) show that the heritability – the measure of relative genetic influences obtained in classic twin studies – of educational attainment is higher in the Nordic countries than in the United States, and also, crucially, that heritability is higher for twins born later as opposed to earlier during the twentieth century (but see Silventoinen et al. 2020). Baier et al. (2022) also show that in Germany, which has a continental welfare regime, and where students are sorted into academic and vocational tracks very early on, the heritability of educational achievement is lower than in countries like Sweden and Norway – countries with social democratic welfare regimes, and with postponed tracking.

Although the above-mentioned studies show contextual differences in heritability, they do not say anything about the effects of genetic propensities in absolute terms. In a study relying on the previously mentioned polygenic index for educational attainment (EA PGI) – a genome-based indicator that measures an individual’s genetic propensities for educational attainment (see further description below) – Lin (2020) finds suggestive evidence of an increasing effect on educational attainment in a sample of Americans born 1920–1959 (cf. Herd et al. 2019). Interestingly, two slightly older studies by Conley et al. (2016) and Okbay et al. (2016) instead shows a decreasing effect of an EA PGI on educational attainment in samples of Americans born 1919– 1955, and Swedish twins born 1929–1958, respectively. The PGI used in each of these studies is based on older genetic findings, however, and mainly include cohorts born during the first half of the twentieth century. Nevertheless, these conflicting results do highlight the need for further investigation.

On the micro level, a socioeconomically favourable family environment may also allow for a complementary interaction to occur. As put by Erola et al. (2022, p. 3), a favourable childhood environment, featuring for example highly educated parents, “can boost the positive genetic influences” on educational attainment. Highly educated parents might for instance task a talented child with extra, appropriately challenging schoolwork. This would be an example of *proximate processes* (Bronfenbrenner and Ceci 1994): continuous environmental stimuli that promotes the development of a given genetic propensity (in this case for education). Inspired by previous twin studies on cognitive ability (e.g. Turkheimer et al. 2003), two recent twin studies support the complementarity hypothesis on the micro level by showing higher heritability of educational attainment for twins with more educated parents (Baier and Lang 2019; Erola et al. 2022). Similar results are also found in recent studies using polygenic indices. Papageorge and Thom (2020) find a stronger association between an EA PGI and college completion for Americans (born in the first half of the twentieth century) with stronger socioeconomic backgrounds (cf. Uchikoshi and Conley 2021). Ronda et al. (2022) corroborate this finding in a sample of Danes born 1985–2005, using various measures of family (dis)advantage. These findings are particularly interesting since they are based on quite recent cohorts growing up in an egalitarian, Scandinavian welfare regime.

Based on previous literature, then, a reasonable expectation would be that favourable environments complement genetic propensities, increasing their influence. One should, however, also discuss the potential for the opposite type of mechanism: where favourable environments instead act as *substitutes* for favourable genetic propensities. As argued by Baier et al. (2022), in a perspective also formulated by (Saunders 2010), an individual with genetic propensities for education that grows up in an unfavourable family environment might persevere precisely *because* of these propensities (a ’challenging’ mechanism). Conversely, to grow up in a socioeconomically favourable environment could mean less of a need to rely on personal abilities in order to reach certain educational goals. True enough, Papageorge and Thom (2020) find a negative interaction between the EA PGI and childhood socioeconomic background when it comes to upper-secondary school completion. Using the same data, Lin (2020) also finds that parental education decreases the effect of the EA PGI on educational attainment. There is therefore reason to remain open with regards to the direction of the interaction, at least in the case of the interaction between genetic propensities and socioeconomic background.

It has so far been argued that both macro- and micro-level environment can be important moderators of genetic influences in their own right, and that a likely (but not certain) mechanism is that of complementarity: favourable environments increase genetic influences on educational attainment. A key argument in this paper, however, is that the macro-level environment, apart from being a potential direct moderator of genetic influences, may also affect the downstream moderating effects of more proximate micro-level environments (Isungset et al. 2022). This is to say that there may be some degree of *interdependence* between these levels, which needs to be taken into account in a model of gene-environment interaction.

Theoretically, it would be possible for the macro-level environment to become more favourable at the same time as there remains a complementary interaction between genetic propensities and socioeconomic background. The benefits of educational expansion, say, might accrue mainly with the well-off in society, leading to a ’Matthew effect’ (Ceci and Papierno 2005; Esping-Andersen 2005). But as argued by Isungset et al. (2022, p. 4), in a discussion of the moderating role of welfare states, ”[i]f welfare state policies influence genetic associations, they would work through intermediate institutions, such as schools, families, and neighborhoods”. An instructive example for comparison is the much-researched ’Scarr-Rowe’ hypothesis, which posits a positive interaction between genetic factors and socioeconomic background in the case of cognitive ability (see e.g. Turkheimer et al. 2003). Importantly, meta-studies have suggested that this interaction is prevalent in the US, but not in western Europe (e.g. Tucker-Drob and Bates 2016). One potential interpretation of this is that the interaction itself may depend on macro-level factors that compensate for socioeconomic background – such as income equality and educational institutions (*ibid.*).

This type of idea is also put forward in the already mentioned study by Lin (2020), which suggests that an initial (negative, i.e. substitutive) interaction between an EA PGI and socioeconomic background, with regards to educational attainment, decreases over time. The argument that will be made here is inspired by this study, but states instead that a more favourable macro-level environment, particularly in terms of equality, serves to decrease or remove any *complementary* interaction between genetic propensities and socioeconomic background. In other words, the added benefit of growing up with family socioeconomic advantage, given existing genetic propensities for education, should decrease as the macro-level environment becomes more favourable, and more equal in particular.

### 2.1 Macro-level developments in twentieth-century Sweden

During the twentieth century, Sweden went from being one of Europe’s poorest countries to being one of its most affluent and egalitarian. Through a combination of rapidly increasing wealth, and a strong social democratic momentum, Sweden developed a comprehensive and universal welfare state via a number of social reforms. Income inequality, as measured by conventional measures like the Gini coefficient, decreased, particularly in the post-war period (Björklund and Palme 2000; Gustafsson and Johansson 2003; Lindbeck 1997).

A series of reforms of the educational system were also implemented. The twenties through the forties saw major expansions of lower-secondary schools, especially in rural areas, lowering the average distance to post-primary education (Lindgren et al. 2019). A milestone was the parliamentary committee, appointed in 1946, whose proposals would include extended compulsory education and postponed tracking, both to the ninth grade, as well as a standardized, national curriculum. An explicit purpose behind these proposals was to increase educational equality of opportunity (Holmlund 2008). Compulsory education had already been partially extended in the fourties (Fischer et al. 2020), but this more comprehensive reform was then gradually rolled out in the fifties (Meghir and Palme 2005). While extended compulsory schooling causes a purely environmental effect on educational attainment, it would indeed be expected to also have opportunity-equalizing effects. True enough, both reforms affected educational attainment particularly for low-SES children. Other reforms implemented during the same period also include subsidized school lunches (Lundborg et al. 2022), and subsidized child care. Later decades would also feature an expansion of higher education, this too with partially egalitarian motives (Gribbe 2022).

### 2.2 Hypotheses

There are, then, a number of important macro-level developments in Sweden during the twentieth century that, taken together, may be expected to have a moderating effect on the relationship between genetic propensities and educational attainment. In line with the complementarity hypothesis, my first hypothesis (H1) states that *the effects of genetic propensities on educational attainment are higher for individuals born late in the twentieth century, compared to individuals born early in the twentieth century.* As for the moderating effect of socioeconomic background, I proceed with the expectation that socioeconomic background also has a positive, *complementary* moderating effect on genetic influences. To this, however, I add an important qualification: that this interaction depends on macro-level context. Thus, my second hypothesis (H2) states that *the effects of genetic propensities on educational attainment are increased by socioeconomic background for individuals born in the early twentieth century, but not for individuals born in the late twentieth century*.

## 3 Data, variables, and methodological approach

This section presents the data, variables, and methodological approach of the study. A data availability and ethical approval statement, and more information about the included variables are included in the appendix. For instance, it includes descriptive statistics for all variables, and an extended presentation of the rationales and procedures behind creating polygenic indices.

### 3.1 Capturing macro-level environments using birth year differences among STR twins

The study uses a long panel of genotyped twin cohorts in the *Swedish Twin Registry* (STR). STR, established in the late 1950’s, is a large and well-known resource for genetically informed research in Sweden (Zagai et al. 2019), in which cohorts of twins are regularly invited to participate. STR contains 43 000 genotyped twins, born between 1911–2005 (see figure 1). The actual birth cohorts studied in the analyses will be restricted, however, by the availability of register data on educational attainment (see below), to which the twins are linked via anonymized personal identifiers. As also shown in the figure, STR contains a number of complete dizygotic twin pairs. These DZ twin pairs will later be used to estimate causally robust within-family analyses, alongside conventional between-family analyses.

**Figure 1:**
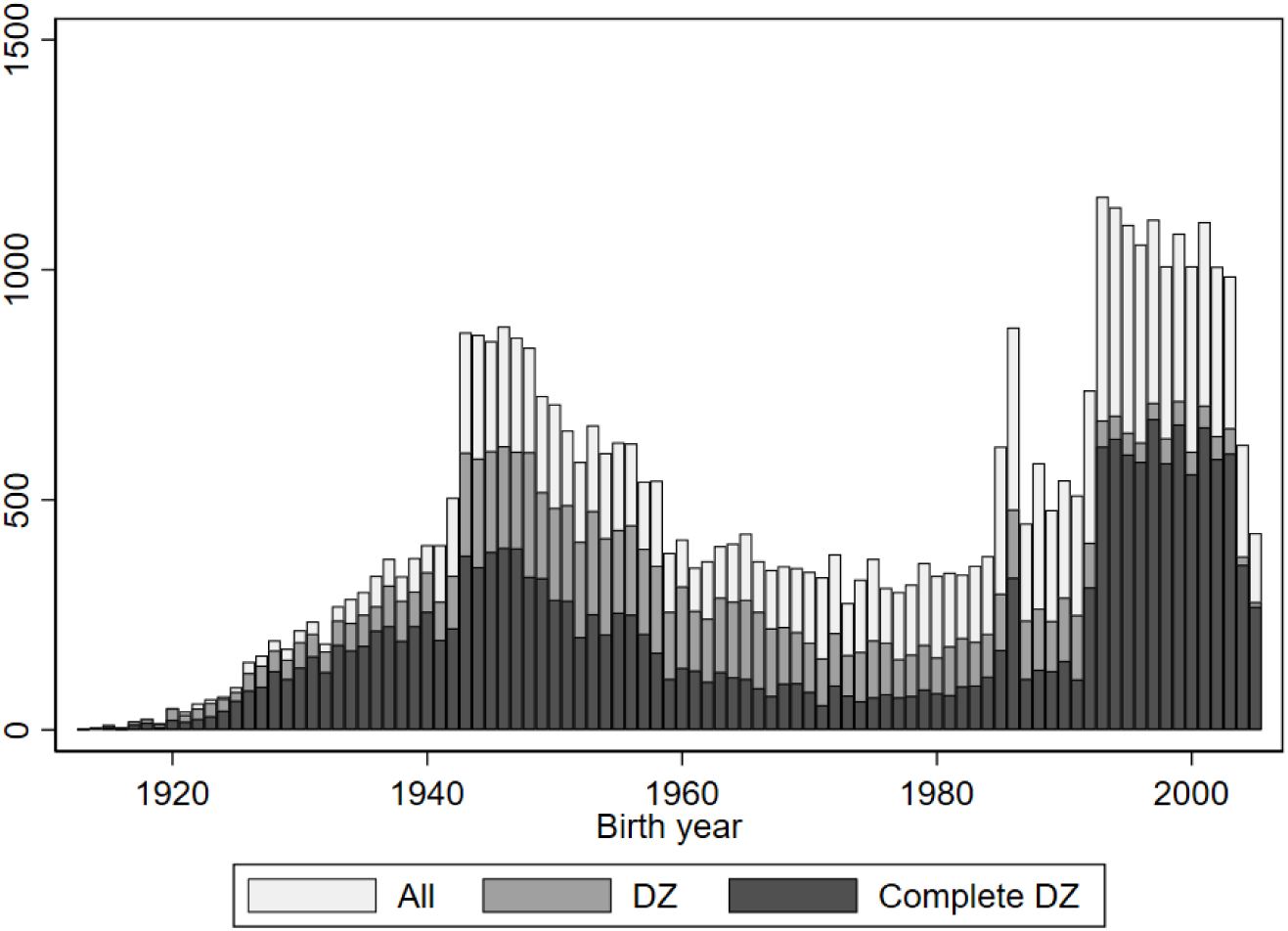
Genotyped twins in STR

Inspired by previous research, the approach is to capture the macro-level moderation of genetic associations with educational attainment by examining whether the association differs depending on when the different STR cohorts are born (e.g. Okbay et al. 2016). The approach has two main advantages. First, as put by Wedow et al., ”[d]ifferences in genetic effects across birth cohorts may be interpreted as reflecting moderation by the *social, institutional and physical conditions* unique to each cohort” (Wedow et al. 2018, p. 806, emphasis added). Analysing birth-year differences should, in other words, be appropriate for testing the hypotheses. Second, a birth cohort naturally cannot be self-selected into, thereby alleviating concerns about endogeneity in the form of gene-environment correlation. The key limitation of the approach, however, is that specific macro-level factors remain ’black-boxed’; discerning the mechanisms through which an interaction with the macro-level environment occurs needs to be based on knowledge about the specific factors that change during the studied period, and which of them should be of key importance.

### 3.2 Twin and parent educational attainment from Swedish national registers

The main dependent variable is educational attainment, measured as a twin’s maximum years of education as counted in the available Swedish population registers (1970, and 1990–2018). Twins that are too young in the latest register are excluded from the analyses, since they will not have had sufficient time to accumulate education. 1989 is accordingly picked as the latest birth cohort in all analyses of educational attainment. The earliest birth cohort in the *main sample*, in which the interaction between genetic propensities and macro-level context is studied, is born in 1925 (the number of genotyped twins in the sample born before 1925 is negligible). The main sample contains in total 28 500 twins, and 11 000 dizygotic twins in full pairs.

Parental education is used to capture socioeconomic background. Although there are alternatives (occupation or income, for example), parental education is commonly used as a proxy for socioeconomic background (Breen and Jonsson 2005), and is also the indicator used in related work (e.g. Lin 2020; Papageorge and Thom 2020). Moreover, parental education can be measured for quite early-born cohorts using the available registers. Specifically, 1940 is set as the earliest birth cohort in the *parental sample*, i.e. the sample where twins can be linked to their parents’ education, and where the (changing) interaction between genetic propensities and socioeconomic background is studied. This sample contains 20 000 twins in total, and 7 500 full-pair DZ twins.

Parental education is measured as the highest education years of either the mother or father, whose respective education is measured in the same way as for the focal twins. This specification minimizes missing data for parental education, but means assuming a gender-uniform influence of parental education. Given the possibility of gendered effects (the mother might on average spend more time with a child during formative years, for instance), supplementary analyses include separate tests for maternal and paternal education.

Since the value of education years in absolute terms has changed substantially during the twentieth century, I decile-rank the educational attainment of the twins, and that of their parents, within each birth cohort, and within gender. Access to full-population register data allow for these deciles to be calculated relative to the equivalent birth cohorts in the full Swedish population, and not just the twin sample. This is likely to improve precision to some extent, since there is a slight over-representation of the highly educated within STR as opposed to the population at large (approximately 0.4 education years on average).

### 3.3 Measuring genetic propensities for educational attainment using a polygenic index

The main independent variable, and that is used to capture the twins’ genetic propensities for educational attainment, is a *polygenic index for educational attainment* (EA PGI). An EA PGI is a individual-level genetic predictor of educational attainment, based on the combined effects of a set of *single-nucleotide polymorphisms* (SNP). SNP’s are locations in the genome where individuals vary on single combinations of AT and CG, and that are associated with educational attainment. SNP’s that are associated with educational attainment are first discovered in a *genome-wide association study* (GWAS). A GWAS runs regressions of millions of different SNP’s on an outcome (here educational attainment) in order to detect whether particular SNP’s are associated with the outcome. The rationale behind GWAS is that most social science outcomes are influenced by a very large number of genetic variants, each with very small effects (Chabris et al. 2015). Expressed formally, a polygenic index for an individual *i* is the sum of minor alleles *x_ij_* at SNP *j*, weighted by the beta coefficient of SNP *j*, as discovered in a preceding GWAS:

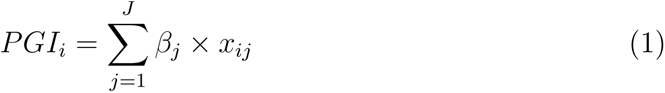

The EA PGI that is constructed for the STR twins is based on the polygenic index repository by Becker et al. (2021), and builds on the latest GWAS for educational attainment (Okbay et al. 2022), which found approximately 4000 SNPs associated with educational attainment in a study of 3 million individuals. The PGI repository allows for two types of EA PGI’s to be constructed: one that is based only on GWAS results for educational attainment (the type described above), and one that also uses GWAS results for a series of supplementary outcomes in order to enhance the predictive accuracy *for* educational attainment. Multi-trait PGIs have been shown to outperform standard, single-trait PGIs in terms of predictive accuracy (Becker et al. 2021; Turley et al. 2018). In order to produce as well-powered analyses as possible, the main analyses will therefore be based on the multi-trait version of the EA PGI. It will however be important to show, as a robustness check, that similar results can be obtained using the single-trait PGI. The EA PGI is standardized by birth cohort in order to avoid potential batch effects (related to the actual genotyping), and the beta coefficient for the EA PGI will therefore show how one standard-deviation increase in the PGI is associated with a one-decile increase in educational attainment.

Having described the EA PGI, it is important to discuss the challenges and limitations that accompany analyses based on polygenic indices, both generally and the EA PGI specifically. A distinction can here be made between methodological and interpretative challenges. Beginning with the methodological dimension, a general limitation is that the GWAS sample on which the PGI is based is not infinitely large. Despite GWAS samples becoming larger and larger (Mills and Rahal 2019), the fact that effects of individual SNP’s are so small means that even huge samples cannot detect all potential genome-wide associations. This introduces attenuation bias in the form of measurement error in the PGI (Biroli et al. 2022).

Another key challenge, which instead biases associations upwards, is *population stratification* (Price et al. 2010). Systematic differences in genetic ancestry between populations and sub-populations, which arise due to non-random mating across social or geographic barriers, may give rise to spurious associations between SNPs and outcomes, if not controlled for. The standard procedure for handling population stratification in analyses of unrelated individuals is to control for a series of principal components of the genetic data.

Receiving increasing attention in the literature is also *genetic nurture*, defined by Trejo and Domingue (2018, p. 188) as the “social genetic effect that parents have on their children.” In short, an individual may be affected by his or her parents’ genotype via environmental pathways. For example, Kong et al. (2018) show that parental genes that are *not* passed on to their children are still associated with their children’s educational attainment. Genetic nurture is expected to inflate EA PGI estimates. By implication, it also means that EA PGI estimates cannot be given a purely genetic interpretation, since it includes some parental-environmental effect as well (for a review, see Wang et al. 2021).

Another potential limitation should also be added regarding the specific approach of studying interactions between the EA PGI and birth cohort. There is a possibility that the association between the EA PGI and educational attainment might increase not because of the macro-level environment changing per se – by becoming more favourable – but rather because the GWAS, on which the EA PGI is based, is computed on a sample that is biased towards younger cohorts. In other words, the EA PGI may be more predictive for younger cohorts for methodological reasons (see e.g. Domingue et al. 2020). At present, there is no way to ascertain whether, or to what extent this challenges the validity of the birth cohort approach, but it must nevertheless be regarded as a potential limitation (although primarily for testing H1).

Turning to the interpretative dimension, it is important to note that the EA PGI is not simply a proxy for educational ability. EA PGI associations capture *any* genetic variant that is (probabilistically) associated with educational attainment (Barcellos et al. 2021), and are expected to be mediated by a mostly unentangleable web of traits, behaviours, and environments. EA PGI associations therefore need to be interpreted cautiously in relation to concepts like meritocracy (Harden et al. 2020), and this is also why, throughout this paper, the more neutral term ’genetic propensities’ has been used instead of more normatively loaded concepts, such as ’genetic endowment’. These caveats remain when the effect of the EA PGI can be causally identified (see below). It may be noted, however, that recent work finds *cognitive* as well as *non-cognitive skills* to partly mediate EA PGI associations with educational outcomes (Malanchini et al. 2023; Rustichini et al. 2022).

### 3.4 Triangulating with between-family and causally robust within-family models

Acknowledging the above-mentioned methodological challenges when it comes to studying PGI associations in a sample of unrelated individuals, this study applies both between-family *and* causally robust within-family models to test its hypotheses. A between-family model, then, uses individual, theoretically unrelated twins as units of analysis, which is also (despite its issues) the most commonly used approach in the social science genomic literature to date. Crucially, and as noted above, this model is expected to be confounded by factors such as population stratification and genetic nurture. A within-family model instead uses the variation that exists between *siblings* in their respective EA PGI and educational attainment. This model allows for a causal interpretation of genetic influences, since the genetic differences between two siblings are outcomes of a true genetic lottery (following Mendel’s second law of independent assortment). By implication, population structure and genetic nurture is expected to be constant within a twin pair. Consequently, this model constitutes the “gold standard to correct for confounds in genetic association studies” (Harden and Koellinger 2020, p. 569). Relying on twins as opposed to normal siblings is also particularly advantageous, since any potential birth order effects are effectively mitigated.

Fitting a within-family model has been shown to reduce EA PGI associations with educational attainment by as much as 60 percent (Selzam et al. 2019) – illustrating the issue of confounding from population stratification and genetic nurture in a model using unrelated individuals. However, using a within-family model with a PGI that is based on a GWAS that has been run *between families* has been shown to introduce additional measurement error (Trejo and Domingue 2018). The within-family estimates are therefore likely to be biased downwards for methodological reasons. As a consequence, gene-environment interactions are also likely to be estimated with less precision. It can also be noted that the amount of variation is expected to be lower between siblings, as opposed to unrelated individuals. Additionally, the number of observations decrease considerably in STR when restricting to full dizygotic twin pairs (see figure 1). In sum, there are a number of factors that affect the precision by which the within-family model can be estimated. For this reason, and as a way of triangulation, results from both between-family and within-family models will be shown.

The respective empirical models will now be described more in detail. In the between-family model, *Edu_ij_* represents the education decile for twin *i* in twin pair *j*. *EAPGI_ij_* is the polygenic index for educational attainment. The Λ*_ij_* term includes controls for the first 20 principal components of the genetic data, and, following convention, interactions between birth year and sex. Standard errors are clustered on the level of twin pair *j*.

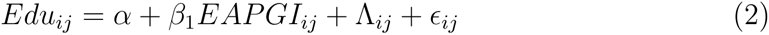

In the within-family model, the object of analysis is instead dizygotic twin pair *j*, and the outcome is defined as the *difference* in educational attainment within a pair, Δ*Edu_j_*. Similarly, the model uses the twin-pair difference in the EA PGI, Δ*EAPGI_j_*. Note that this within-pair differencing is only performed for the outcome and the EA PGI. Parental education and birth year remains measured *between* twin pairs. The Ω*_i_j* term is a control for sex difference within twin pairs.

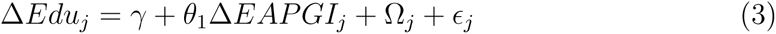

## 4 Results

Does the relationship between the EA PGI and educational attainment become stronger over time for the Swedish twin cohorts? Figure 2 shows the estimates for equation 1 (between-family) and 2 (within-family) in birth periods spanning 1925-1949, 1950-1969, and 1970-1989. The average association is almost twice as high in the between-family model compared to the within-family model (0.7 deciles, and 0.4 deciles, respectively). As explained above, the between-family association is expected to be biased upwards due to confounding. The within-family association should not be confounded by factors like population stratification and genetic nurture, but is also expected to be attenuated due to measurement error. More importantly, though, the figure illustrates that the associations, using both models, do indeed increase with birth period: the associations increase from the oldest birth period (1925–1949) to the youngest birth period (1970–1989). Specifically the association increases by approximately 27 percent in the between-family analysis, and 43 percent in the within-family analysis.

**Figure 2:**
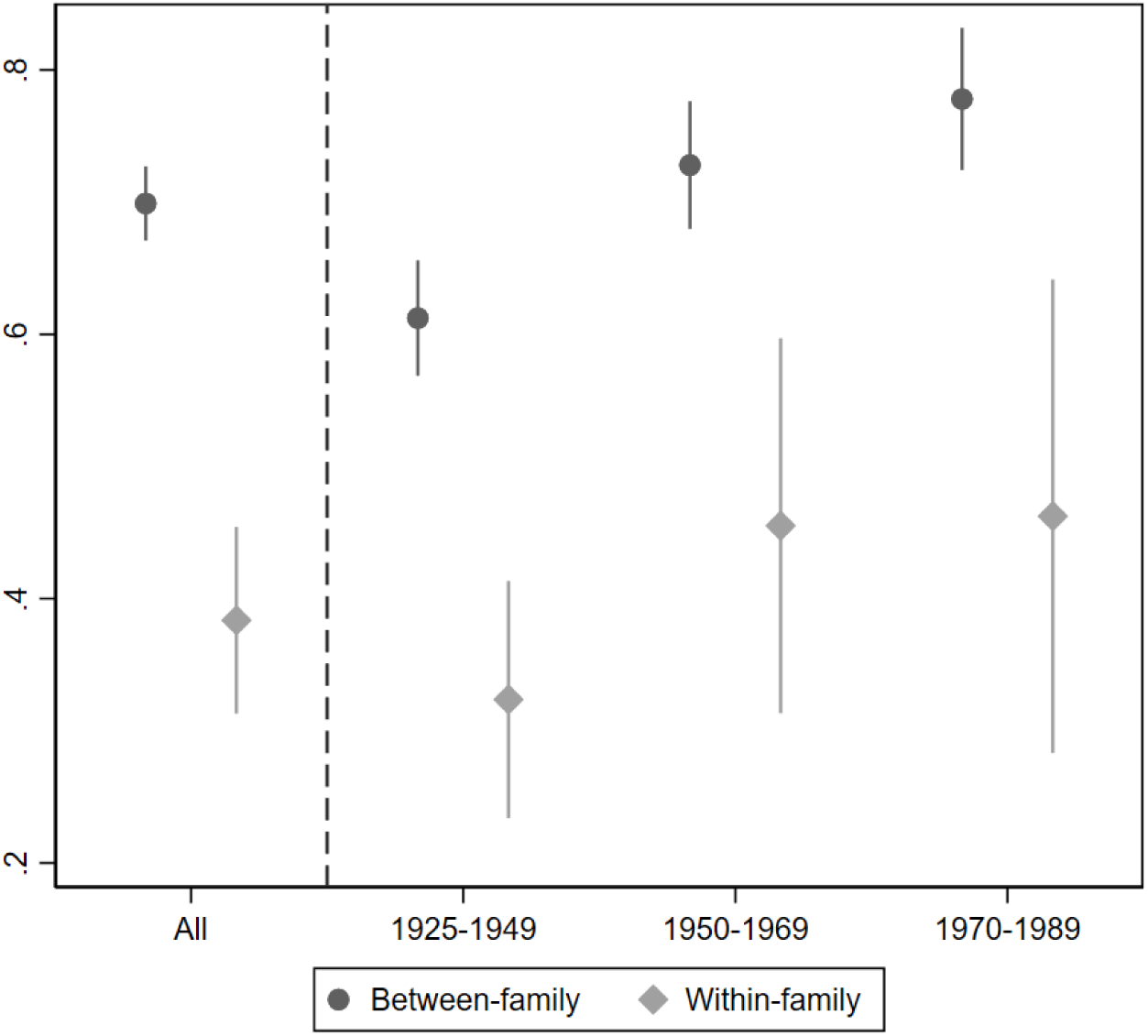
Relationship between EA PGI and educational attainment by birth period Note: The Y-axis measures the EA PGI coefficient, across all birth cohorts, and separately for three different birth periods: 1925–1949, 1950–1969, and 1970–1989. Estimates from the between-family model includes controls for the first 20 principal components of the genetic data, and an interaction between birth year and sex. The within-family model includes a control for sex difference within a twin pair. Estimates are shown with 95 percent confidence intervals.

To test whether these increases are statistically significant, two types of interaction models are estimated (see ’Equations’ in the appendix, and tables A4–A5). The first one corresponds directly to figure 2, and uses dummies for the birth periods 1950– 1969, and 1970–1989, with 1925–1949 as the reference category. The between-family differences between the oldest birth period and each of the two younger birth periods are statistically significant (*p*=0.001, and *p*=0.000). An F-test of the joint significance of the interaction terms is also statistically significant (*p*=0.000). While the increases in the PGI coefficient are roughly the same in the within-family model, they are not strictly significant (*p*=0.140, and *p*=0.168), as is also suggested by the coefficient plot. Based on the discussion above, however, these noisier results from using the within-family model is not unexpected.

The second interaction model estimates a linear interaction, using a continuous birth year variable. This model provides statistically significant interaction estimates of 0.004 (*p*=0.000) in the between-family model, and a statistically significant 0.005 (*p*=0.032) in the within-family model. The association between the EA PGI and educational attainment thus becomes 0.004/0.005 deciles larger per birth year, which in turn translates to approximately a quarter decile across the entire panel. Using the single-trait EA PGI instead of the multi-trait EA PGI provides very similar results (see figure A2 and tables A8–A9).

I now turn to the question of whether the association between the EA PGI and educational attainment is positively moderated by parental education, and crucially, whether this interaction differs depending on period of birth. Figure 3, with the baseline birth year now instead being 1940, illustrates the association when each sub-sample is split by median parental education. The figure suggests a slight positive interaction between genetic propensities and parental education when averaging across all birth cohorts. In line with hypothesis 2, however, the interaction appears to depend on birth year. The interaction is at its strongest in the cohorts born 1940– 1949, and decreases in the subsequent birth periods. The changes are most apparent from looking at the within-family model, but the pattern is the same for both models: there is an initial positive moderation of parental education that tapers off over time, nearing zero, and even turning negative – although it cannot clearly be concluded that these interactions are statistically different from zero.

**Figure 3:**
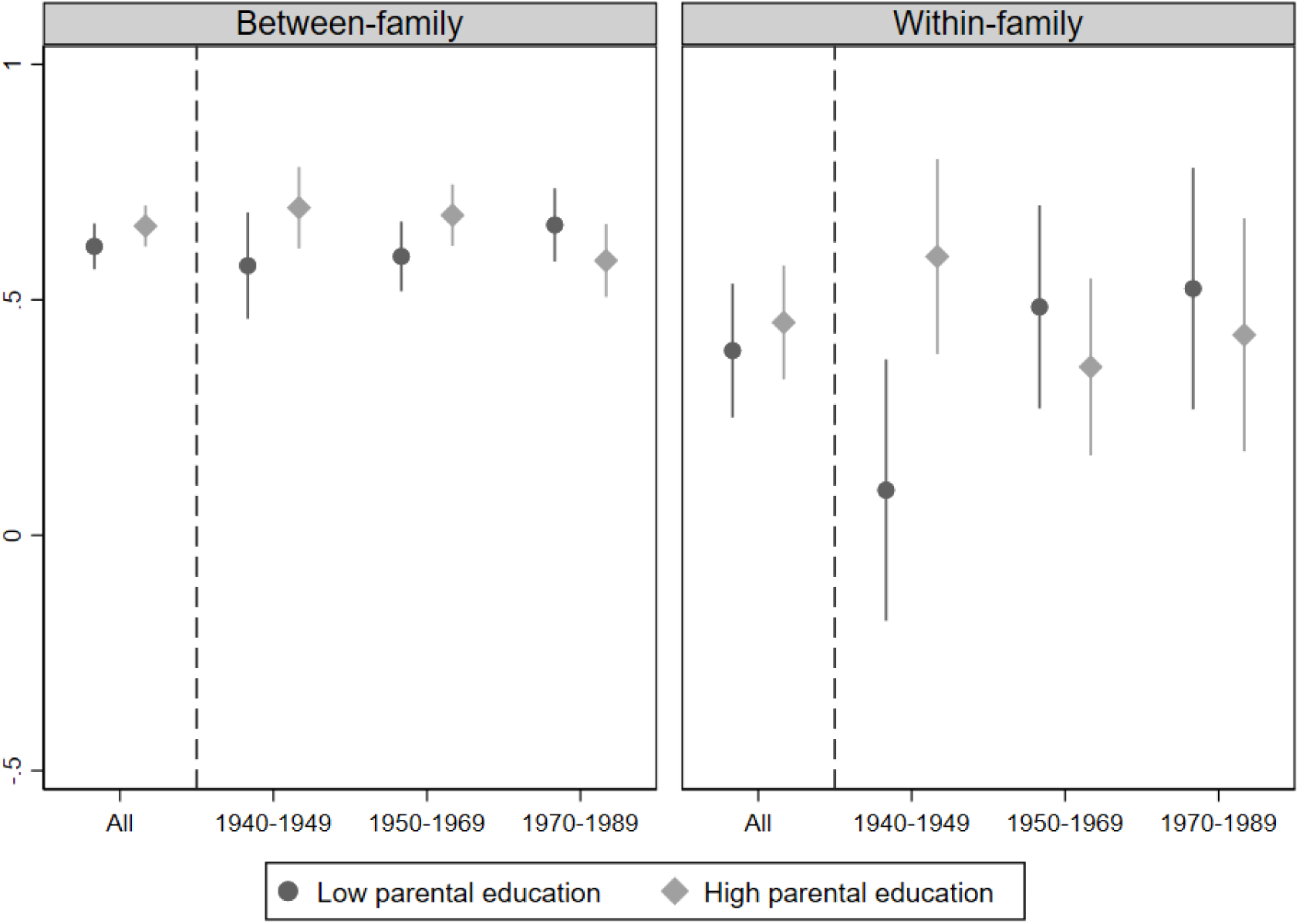
Relationship between EA PGI and educational attainment by birth period, split by parental education Note: The Y-axis measures the EA PGI coefficient, across all birth cohorts, and separately for three different birth periods: 1940–1949, 1950–1969, and 1970–1989, split by median parental education. Estimates from the between-family model includes controls for the first 20 principal components of the genetic data, and an interaction between birth year and sex. The within-family model includes a control for sex difference within a twin pair. Estimates are shown with 95 percent confidence intervals.

Three-way interactions are now calculated using the EA PGI, parental education, and birth period/year (see ’Equations’ in appendix, and tables A6–A7). The birth period model gives that the interaction between genetic propensities and parental education decreases in both the between-family and within-family model. For the between-family model, the difference between the oldest birth period (1940-1949) and the middle birth period is barely significant at the 0.1-level (*p*=0.116), whereas that between the oldest group and the youngest group is significant (*p*=0.014). The F-test of joint significance is statistically significant (*p*=0.028), however. The equivalent within-family differences are both significant (*p*=0.004, *p*=0.005), and the F-test is also significant at *p*=0.013).

The continuous birth year model provides a significant negative three-way interaction coefficient for the between-family model (*p*=0.023). The interaction coefficient for the within-family model is, however, not significant on any conventional level (*p*=0.232). As in the previous analysis, the results from using the single-trait EA PGI are very similar (see figure A3 and tables A10–A11). As another graphical illustration of how the interaction between the EA PGI and parental education evolves over time, figure A15 shows rolling regressions of the interaction with parental education. This shows how the linear interaction coefficient is initially positive in both models, then decreases quite rapidly, and hovers around zero for all twin cohorts born approximately after 1950.

### 4.1 Supplementary analyses

Supplementing the main analysis, an analysis is made that includes twin cohorts too young to be included in the main sample (born up to 1998). Here, the dependent variable is instead a dummy for having any amount of *tertiary education*, i.e. education beyond upper-secondary school. The results from this analysis corroborate the main results, with a generally increasing association between the EA PGI and tertiary education, and a decreasing interaction between the EA PGI and parental education (see figure A8–A9 and A12–A15). Analyses were also made using an indicator for *upper-secondary school performance* (GPA), for which data is available for twin cohorts born 1955–2000. Although this sample does not include cohorts as old as those in the main analyses, the results for upper-secondary school performance strongly echo those for educational attainment (see figure A10–A11 and tables A16– A19). A different, yet important indicator of status is *income*, which can be measured in early adulthood for twins born 1940–1989. Interestingly, there is no clear evidence of an increasing effect over time on income. The interaction with parental education, however, follows a similar pattern as for educational attainment (see figure A12–A13 and tables A20–A23). The results from the main analyses can therefore be shown to mostly extend to a number of related, yet potentially distinct educational and economic outcomes.

The results regarding the interaction between the EA PGI and parental education were also re-estimated separately for maternal and paternal education. Results do not change markedly depending on parental gender, suggesting that the assumption of gender-uniform moderating effects of parental education appears was appropriate (see figures A5–A4).

Finally, the between-family results are re-estimated using only the full dizygotic twin pairs in the sample. The between-family model in the main analysis included dizygotic as well as monozygotic twins in order to maximize the number of observations. It has also included twins whose sibling is not genotyped, or not present in the data. Whereas in the main analysis the sample restriction differs between the between-family and within-family model, in these robustness analyses the exact same DZ twin sample is used for both models. Making this restriction for the between-family model (the within-family model is not affected, since it only uses full DZ pairs), the increase in the main effect over time becomes slightly weaker, but the overall patterns appear robust (see figures A6–A7).

## 5 Discussion

Combining conventional between-family analyses with causally robust within-family analyses, the results suggest that the association between Swedish twins’ genetic propensities for educational attainment and actual educational attainment depend on birth period, so that the association is stronger for twins born in the late twentieth century than for those born in the early twentieth century. This provides support for hypothesis 1. In terms of the paper’s theoretical framework, the twins grow up in substantially different macro-level institutional and sociopolitical environments, with the later-born twins growing up in a more favourable environment – one with a fully established, and education-centered welfare state – which in turn increases the influence of their genetic propensities for educational attainment. Moreover, and in support of hypothesis 2, socioeconomic background has also been shown to moderate the association in a complementary manner, but only for the early-born twins – for whom the main compensatory functions of the welfare state were not yet as developed. No existing study has used as an extensive sample of birth cohorts to explore macro-level gene-environment interactions in a single country over time. The results echo recent studies (Conley et al. 2016; Herd et al. 2019; Lin 2020; Okbay et al. 2016), but extend well beyond them in terms of its scope. The results also echo Branigan et al. (2013), who finds an increasing heritability of educational attainment over time. It has remained unsettled in the twin study literature whether the heritability of educational attainment has increased due to genetic factors having higher effects, or by becoming relatively more important due to diminishing effects of the environment (Engzell and Tropf 2019). This study suggests that the association between genetic propensities and education have gotten stronger over time in *absolute* terms, at least in Sweden.

The results concerning the interaction between genetic propensities and socioeconomic background corroborate some previous studies, but, with the novelty of showing how the interaction evolves over time, quite literally puts them into context. Indeed, the positive interaction found in e.g. Papageorge and Thom (2020) is based on a sample of American birth cohorts similar in time to the oldest cohorts in this study. The interaction found there might therefore, just as the one found here, be temporally dependent. The null-findings for the later-born cohorts do however contradict Ronda et al. (2022), who found a positive interaction in a Danish sample using rather young cohorts. One might have expected similar findings, given the countries’ sociopolitical and institutional similarities. However, while the twins in STR are generally slightly more educated than the population, the sample used by Ronda et al. (2022) investigates the ”genetic and environmental architecture of severe mental disorders” (p. 7). The STR sample should therefore arguably be more representative of the population. This study should then illustrate the importance of recognizing macro-level environments when studying both the association between genetic propensities and educational attainment per se, *and* the interaction between genetic propensities and socioeconomic background with regards to educational attainment. Given that both might differ according to the birth cohorts used, generalizations from single studies must be done with caution (see e.g. Isungset et al. 2022).

While having several advantages, this study is not free from limitations. First, any moderating effect of birth year is clearly a due to a combination of factors, the relative importance of each is difficult to determine. It should however be reasonable to argue that it is primarily the developments in educational institutions and economic inequality that are most crucial, and that drive interactions with genetic propensities – particularly so during the specific period under study. Second, there is the possibility that the stronger association for the younger cohorts comes about because of a better fit, and not because of gene-environment interaction. However, and going against this alternative interpretation, one would then also have expected to be less likely to find significant interactions with socioeconomic background in the oldest cohorts. One could also argue that if the stronger association comes about only due to better fit, the results should have been similar for income (which they are not). Nevertheless, this point should be important to investigate further in future work.

One should also be humble regarding what the association between the EA PGI and educational attainment actually captures. Although recent research suggests that cognitive and non-cognitive skills may well mediate the association, any interpretation related to meritocracy or equality of opportunity is vulnerable to the presence of non-meritocratic mechanisms. If one accepts the results in this study as evidence of increased genetic effects over time due to a more open and favourable environment, however, one reasonable conclusion could be that at least the *increase* in the association can be interpreted in these terms. In other words, understanding the changes in the environment may be key to interpreting these gene-environment interactions in relation to equality of opportunity.

Even so, it remains philosophically ambiguous whether increased genetic effects on educational attainment, even if evenly distributed across socioeconomic background, are preferable. After all, genes are as much out of a person’s control as his or her initial social circumstances. As colorfully put by Sandel (2021, p. 123), ”[i]f our talents are gifts for which we are indebted – whether to the genetic lottery or to God – then it is a mistake and a conceit to assume we deserve the benefits that flow from them.” Still, a core aspect of contemporary understandings of equality of opportunity is that of increasing effects of the individual’s own traits and abilities (e.g. Esping-Andersen and Cimentada 2018). Accordingly, then, this study might indicate a development towards more equal opportunity in Sweden during the twentieth century. The fact of abilities being at some level undeserved does, however, serve as a foundation for a continuing commitment to social equality.

## Notes

### Competing Interest Statement

The authors have declared no competing interest.

