## Supplementary material for "Genetic Influences on Educational Attainment Through the Lens of the Evolving Swedish Welfare State: A cross-level gene-environment interaction study based on polygenic indices and longitudinal register data"

October 18, 2023

### 1 Data overview

#### 1.1 Data availability and ethics approval

The study uses individual-level data from the Swedish Twin Registry (STR), which is administered by the Steering Committee of the Swedish Twin Registry. The data material is located on an encrypted server on to which one has to log in through a remote desktop application in order to perform all of the data analyses. Due to the sensitivity of the data, the author is under contractual and ethical obligation not to distribute these data to others. For those researchers who want to replicate the results, they must obtain approval from the Swedish Ethical Review Authority and from the Steering Committee of the Swedish Twin Registry. Researchers using STR data are required to follow the terms of a number of clauses designed to ensure protection of privacy and compliance with relevant laws. For further information, visit `https://ki.se/en/research/swedish0twin-registry-for-researchers`.

This research has been approved by the Swedish Ethical Review Authority (*Etikprövningsmyndigheten*), [XXXXXX], and by the Steering Committee of the Swedish Twin Registry (*STR:s expertgrupp*).

#### 1.2 Variable sources

Table A1 provides an overview of all the main and supplementary variable used in the study. When applicable, it shows which original variables from different register sources are combined to create one single variable for a particular outcome. Lastly, it indicates whether a variable is featured only in the appendix, the main text, or in both.

Table A1: Register sources for all variables

| Variable used | Original variable | Data source | Appendix only |
|---|---|---|---|
| Multi-trait EA PGI | PGI_EA_multi | Swedish Twin Registry (STR) | No |
| Single-trait EA PGI | PGI_EA_single | Swedish Twin Registry (STR) | Yes |
| PC1-PC20 | PC1-PC20 | Swedish Twin Registry (STR) | No |
| Years of education | SUN_2000niva | Longitudinal integrated database for health insurance and labour market studies (LISA) | No |
| | UtbNiva | Population and Housing Census (FoB) | |
| Tertiary education | SUN_2000niva | Longitudinal integrated database for health insurance and labour market studies (LISA) | Yes |
| Upper-secondary school performance | mbetyg | Registret över slutbetyg från gymnasieskolan | Yes |
| | jmftal | Registret över slutbetyg från gymnasieskolan | |
| Parental years of education | SUN_2000niva | Longitudinal integrated database for health insurance and labour market studies (LISA) | No |
| | UtbNiva | Population and Housing Census (FoB) | |
| Mother's years of education | SUN_2000niva | Longitudinal integrated database for health insurance and labour market studies (LISA) | Yes |
| | UtbNiva | Population and Housing Census (FoB) | |
| Father's years of education | SUN_2000niva | Longitudinal integrated database for health insurance and labour market studies (LISA) | Yes |
| | UtbNiva | Population and Housing Census (FoB) | |
| Income in early-middle adulthood | ForvErs | Longitudinal integrated database for health insurance and labour market studies (LISA) | Yes |
| | ArbInk | Population and Housing Census (FoB) | |
| | ArbInk | Population and Housing Census (FoB) | |
| Birth year | Birth year | Multigenerational Register (FlerGen) | No |
| Sex | Kon | Multigenerational Register (FlerGen) | No |

#### 2 Presentation of GWAS and polygenic indices

This section provides a primer on genomics, and how genomics may be applied in a social science setting, using polygenic scores. The human genome consists of around 3 billion pairs of nucleotide molecules: adenine, guanine, cytosine, and thymine. Particular stretches of the genome make up *genes*, of which we have 20 000–25 000. Genes are instruction codes for building chains of amino acids (proteins) that regulate how the cell (and therefore the entire organism) functions. Individuals share 99.9 percent of their genomes. The locations in the genome where humans do differ from each other (the remaining 0.1 percent) are called *polymorphisms.* The most common polymorphisms are *single-nucleotide* polymorphisms (SNP), locations in the DNA sequence where there is variation in single nucleotide molecules, i.e. A, C, T or G. Genes can contain hundreds or more SNPs. But, and as is the case in some genetic diseases, for example, a single-DNA letter mutation in a gene can be enough for it to produce partly or entirely dysfunctional proteins. Single-DNA letter differences, then, may be consequential for different *phenotypes.* For most SNPs, only two possible nucleotides occur in the population: the 'major allele', and the 'minor allele'. We all receive one allele, major or minor, from each of our parents, and it follows that we end up having 0, 1, or 2 minor alleles at a given SNP. We count the minor alleles for SNP $j$ to get an individual's *genotype* at SNP $j$.

Probably the single most important genetic insight of the last decade or so is that most individual-level characteristics, traits, behaviours and outcomes are immensely *complex* from a genetic point of view. As such, they are influenced by a very large number of SNPs with small to miniscule effects (Chabris et al. 2015). This insight about the *polygenicity* of complex human traits and outcomes has caused a paradigm shift within the field: from thinking in terms of single genes' effects – as one would have done in the so-called 'Candidate Gene era' (e.g. Caspi et al. 2003) – to thinking in terms of the total effect of the genome. The tool used to discover the different SNPs associated with some outcome of interest is called a 'GWAS', or *genome-wide association study.* A GWAS is a large-scale data analysis in which one tests for statistical associations between millions of SNPs and an outcome, one at a time. The outcome could be e.g. height, BMI. depression, self-rated well-being, or in the present case, educational attainment (Okbay et al. 2022). An association between any one SNP and the outcome is deemed statistically significant only below the stringent p-value of $p <$ 5 x $10^{-8}$. This extreme threshold is adopted in order to reduce the risk for false positives due to multiple testing (Biroli et al. 2022). Worth emphasising is that what is crucial in a GWAS is merely that a SNP turns out to be associated with the outcome in question. GWAS is a hypothesis-free, data-driven approach to mapping associations between SNPs and an outcome. The final output of a GWAS is a long list of SNPs and their corresponding beta coefficients related to the outcome in question.

One particular extension of GWAS has proven to be quite valuable for social science: the construction of *polygenic indices* (PGI). A PGI is a summary index based on the regressions performed in a GWAS. The rationale behind a PGI is, indeed, that it is not the effect of single SNPs that are interesting, but their combined effect. To create a PGI, one needs to have first performed a GWAS on an outcome. With the GWAS in hand, one takes each of the hundreds or thousands of beta coefficients from the GWAS regressions, multiplies them with the number of minor alleles that an individual (contained in another, independent sample) has of each SNP, finally summing them up. Say, as a stylized example, that a person's genome consists of 2 SNPs. A GWAS finds that a SNP version (allele) on the first SNP locus has an effect of 0.5 on the outcome, and one with an effect of 0.3 at the other (numbers imagined). It then turns out that this person has zero copies of the effect allele at the first SNP, but two of it at the other SNP (the individual is *homozygous* for both SNP). The PGI would consequently add up to $(0.5 \times 0)+(0.3 \times 2)=0.6$. Extrapolating to the entirety of the measured genome, a polygenic index for an individual $i$ is the sum of minor alleles $x_{ij}$ at SNP

$j$, weighted by the beta coefficient of SNP $j$:

$$PGI_i = \sum_{j=1}^{J} \beta_j * x_{ij} \tag{1}$$

Having been constructed in an independent sample (that was not used in the preceding GWAS), a PGI can then, finally, be added into a regression as a regular individual-level independent variable. Adding a PGI to a regression produces a single beta coefficient, showing how a one unit increase in the PGI, by convention a standard deviation from the sample mean, is associated with a one unit increase in the dependent variable. A PGI can therefore estimate an attenuated version of the absolute effect of the genome. A PGI can also estimate relative effects; this would be the $R^2$ of the PGI model. Distinguishing absolute effects from relative effects can be important when comparing PGI results with results from classic twin studies.

In the main analyses, a so-called *multi-trait* polygenic index for educational attainment (EA PGI) is used, created from the polygenic index repository by Becker et al. (2021). A single-trait EA PGI is constructed just in the way described above, running a GWAS on a trait/outcome (here educational attainment) and then creating polygenic scores based on the GWAS information. A multi-trait PGI is instead complemented by a series of other, related outcomes in the GWAS stage, in order to enhance the predictive accuracy of the EA PGI for educational attainment. The technique used is called 'Multi-Trait Analysis of GWAS' (MTAG), and is explained in detail in Turley et al. (2018). With MTAG, one essentially leverages the fact that GWAS estimates for traits/outcomes that are related to each other will be to varying extent genetically correlated with each other. The GWAS information for one or more related traits can therefore be used to enhance the predictivness for the focal trait/outcome. While the assumptions underlying MTAG are often violated, multi-trait estimation consistently outperforms single-trait estimation, as noted by Turley et al. (2018). Hence, the main analyses in this paper uses a multi-trait EA PGI, that is then based on multi-trait analysis of GWAS, or MTAG. The specific related traits used to construct the multi-trait EA PGI are (Becker et al. 2021): 1) cognitive ability, 2) math ability, 3) age of first born child, 4) delay discounting, and 5) religious attendance).

### 3 Descriptive statistics and further variable information

#### 3.1 Descriptive statistics for main variables

Table A2 provides descriptive statistics for the main variables (used in the main text). The statistics are shown both for the full sample, and for full DZ pair twins only. In figure A1, and following Papageorge and Thom (2020), I also show the distribution of the multi-trait EA PGI, split by parental education and birth period. These plots show that it is not the very tails of either distribution that are being compared when the EA PGI is interacted with parental education.

Table A2: Descriptive statistics (main variables)

a) Main sample (1925-1989)

| | All | | | | | DZ | | | | |
|---|---|---|---|---|---|---|---|---|---|---|
| VARIABLES | N | mean | sd | min | max | N | mean | sd | min | max |
| EA PGI (multi) | 28,628 | 3.57e-09 | 0.999 | -3.822 | 4.320 | 10,958 | 0.00909 | 0.999 | -3.714 | 4.008 |
| EA PGI (single) | 28,628 | -3.04e-09 | 0.999 | -4.245 | 4.663 | 10,958 | 0.0175 | 0.996 | -3.985 | 4.663 |
| Education years | 29,399 | 12.51 | 2.641 | 7 | 20 | 11,127 | 12.17 | 2.698 | 7 | 20 |
| Birth year | 29,417 | 1,958 | 17.05 | 1,925 | 1,989 | 11,130 | 1,953 | 16.78 | 1,925 | 1,989 |
| Sex (female) | 29,417 | 0.558 | 0.497 | 0 | 1 | 11,130 | 0.553 | 0.497 | 0 | 1 |

b) Parental sample (1940-1989)

| | All | | | | | DZ | | | | |
|---|---|---|---|---|---|---|---|---|---|---|
| VARIABLES | N | mean | sd | min | max | N | mean | sd | min | max |
| EA PGI (multi) | 24,943 | 4.97e-09 | 0.999 | -3.822 | 4.320 | 8,650 | 0.00255 | 0.998 | -3.714 | 4.008 |
| EA PGI (single) | 24,943 | -2.19e-09 | 0.999 | -4.245 | 4.663 | 8,650 | 0.0143 | 0.996 | -3.985 | 4.663 |
| Education years | 25,660 | 12.75 | 2.534 | 7 | 20 | 8,815 | 12.51 | 2.568 | 7 | 20 |
| Parent edu. years | 20,020 | 11.21 | 2.941 | 7 | 20 | 7,444 | 10.98 | 2.996 | 7 | 20 |
| Mother edu. years | 20,020 | 10.46 | 2.666 | 7 | 20 | 7,444 | 10.26 | 2.673 | 7 | 20 |
| Father edu. years | 18,006 | 10.70 | 2.829 | 7 | 20 | 6,501 | 10.59 | 2.893 | 7 | 20 |
| Birth year | 25,674 | 1,962 | 15.11 | 1,940 | 1,989 | 8,816 | 1,959 | 14.79 | 1,940 | 1,989 |
| Sex (female) | 25,674 | 0.570 | 0.495 | 0 | 1 | 8,816 | 0.560 | 0.496 | 0 | 1 |

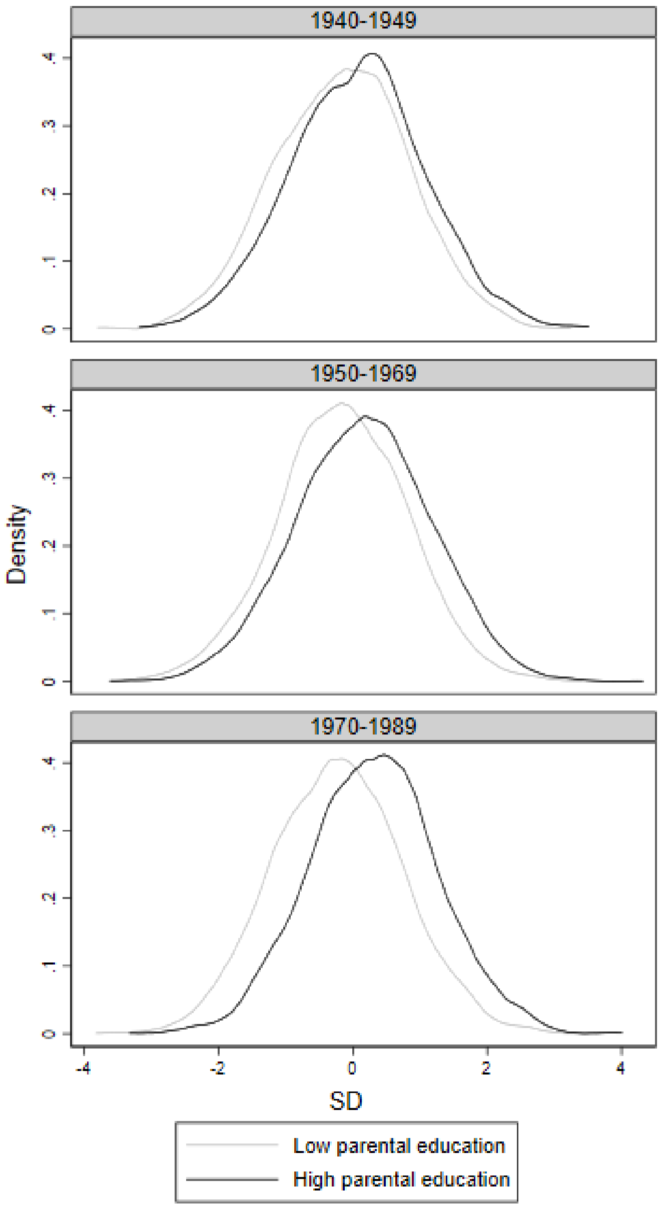

Figure A1: Distribution of EA PGI by birth period, split by median parental education

#### 3.2 Presentation of, and descriptive statistics for supplementary dependent variables

*Tertiary education* is used as a supplementary dependent variable. A twin is measured as having tertiary education (1), as opposed to no tertiary education (0), if he or she is registered as having equal to or more than 13 years of education. An advantage of using this variable is that it allows for an analysis of the interactions with parental education for the youngest possible twin cohorts, specifically those born between 1990–1998. However, it is worth noting that, compared to education year deciles, the results for this variable will be subject to lack of comparability over time. Indeed, the share of individuals receiving tertiary education has increased substantially over time during the studied period. Nevertheless, it is worthwhile studying it as a robustness check with regards to the main results.

*Upper-secondary school performance*, measured in terms of grade point average at the end of upper-secondary school, is also used as a supplementary dependent variable. The purpose of testing this variable is to see whether the results for educational attainment are generalizable to adjacent but potentially different outcomes (in terms of how the effects of genetic propensities are moderated by the environment). Unfortunately, grades data are only available for the 1955 cohort and onwards, meaning that comparisons cannot be made with the oldest cohorts available in the analysis of educational attainment (born 1940–1949). However, the data extend as far forward as to the 2000 cohort, meaning that results for very late-born cohorts can be studied. Additionally, the panel is likely to be long enough to at least detect patterns over time that are substantially different from those in the main analysis, were these to exist.

To account for the fact that the grading system in Sweden has been subject to reform over the years, as well as potential grade inflation, GPA's are also decile-ranked within birth cohorts, relative to the full population. Finally, if the same methodological approach – including birth period dummies as well as a continuous birth year variable, is to be employed for the performance variable, new birth period dummies are required. The new dummies are constructed so as to correspond to the following birth periods: 1955–1969, 1970–1989, and 1990–2000. The oldest birth period in this case therefore corresponds roughly to the middle one in the main analysis.

*Income in early-middle adulthood* is also used as a supplementary dependent variable. Whereas the mechanisms by which individual-level income are determined may not be the very same as for educational attainment, it is still worthwhile to assess whether the patterns for the two variables are similar. Unlike upper-secondary school performance, income can be captured for the same sample used for educational attainment in the parental education analysis, born 1940–1989.

Ideally, one would like to measure income at the exact same time for each cohort. This is, however, not possible due to a limited number of censuses that contain individual-level income data. Measuring income for all cohorts therefore means taking a less restrictive approach to the specific age at which income is measured. 35 is here picked as the 'preferred' age for income measurement, although there will be some variation around this age depending on birth cohort. Income for cohorts born 1940–1944 is measured in 1975 (age 35–31). Income for cohorts born 1945–1954 is measured in 1985 (age 40–31). Income for cohorts born 1955–1982 is measured at age 35, using registers for the years 1990–2017. The latest available register, for the year 2018, is used to measure income for the youngest possible cohorts, born 1983–1989 (age 35–28).

Table A3 provides descriptive statistics for the supplementary variables (used in the appendix).

Table A3: Descriptive statistics (supplementary variables)

a) Income sample (1940-1989)

| | All | | | | | DZ | | | | |
|---|---|---|---|---|---|---|---|---|---|---|
| VARIABLES | N | mean | sd | min | max | N | mean | sd | min | max |
| EA PGI (multi) | 24,943 | 4.97e-09 | 0.999 | -3.822 | 4.320 | 8,650 | 0.00255 | 0.998 | -3.714 | 4.008 |
| Parent edu. years | 20,020 | 11.21 | 2.941 | 7 | 20 | 7,444 | 10.98 | 2.996 | 7 | 20 |
| Income | 25,476 | 1,867 | 1,575 | 0 | 22,364 | 8,765 | 1,634 | 1,461 | 0 | 15,975 |
| Birth year | 25,674 | 1,962 | 15.11 | 1,940 | 1,989 | 8,816 | 1,959 | 14.79 | 1,940 | 1,989 |
| Sex (female) | 25,674 | 0.570 | 0.495 | 0 | 1 | 8,816 | 0.560 | 0.496 | 0 | 1 |

b) Tertiary education sample (1925-1998)

| | All | | | | | DZ | | | | |
|---|---|---|---|---|---|---|---|---|---|---|
| VARIABLES | N | mean | sd | min | max | N | mean | sd | min | max |
| EA PGI (multi) | 36,984 | 4.06e-09 | 0.999 | -3.822 | 4.320 | 15,215 | 0.0124 | 0.997 | -3.714 | 4.008 |
| Tertiary edu. | 38,080 | 0.432 | 0.495 | 0 | 1 | 15,554 | 0.393 | 0.489 | 0 | 1 |
| Birth year | 38,156 | 1,967 | 21.41 | 1,925 | 1,998 | 15,584 | 1,965 | 23.57 | 1,925 | 1,998 |
| Sex (female) | 38,156 | 0.550 | 0.498 | 0 | 1 | 15,584 | 0.534 | 0.499 | 0 | 1 |

c) Tertiary education sample (1940-1998)

| | All | | | | | DZ | | | | |
|---|---|---|---|---|---|---|---|---|---|---|
| VARIABLES | N | mean | sd | min | max | N | mean | sd | min | max |
| EA PGI (multi) | 33,299 | 5.17e-09 | 0.999 | -3.822 | 4.320 | 12,907 | 0.00862 | 0.997 | -3.714 | 4.008 |
| Tertiary edu | 34,341 | 0.454 | 0.498 | 0 | 1 | 13,242 | 0.420 | 0.494 | 0 | 1 |
| Parent edu. years | 26,874 | 11.84 | 2.972 | 7 | 20 | 11,783 | 11.99 | 3.038 | 7 | 20 |
| Birth year | 34,413 | 1,970 | 19.35 | 1,940 | 1,998 | 13,270 | 1,971 | 21.02 | 1,940 | 1,998 |
| Sex (female) | 34,413 | 0.557 | 0.497 | 0 | 1 | 13,270 | 0.535 | 0.499 | 0 | 1 |

d) Upper-secondary performance sample (1955-2000)

| | All | | | | | DZ | | | | |
|---|---|---|---|---|---|---|---|---|---|---|
| VARIABLES | N | mean | sd | min | max | N | mean | sd | min | max |
| EA PGI (multi) | 25,016 | 3.61e-09 | 0.999 | -3.822 | 4.320 | 9,653 | 0.0393 | 0.991 | -3.631 | 4.008 |
| GPA pre-1997 | 7,533 | 338.0 | 61.43 | 75 | 500 | 2,141 | 337.9 | 62.31 | 114 | 494 |
| GPA post-1997 | 14,596 | 14.79 | 3.733 | 0.600 | 310 | 6,467 | 14.75 | 2.789 | 0.600 | 20 |
| Parent edu. years | 21,361 | 12.78 | 2.725 | 7 | 20 | 9,770 | 13.09 | 2.681 | 7 | 20 |
| Birth year | 26,017 | 1,982 | 14.41 | 1,955 | 2,000 | 10,028 | 1,985 | 14.61 | 1,955 | 2,000 |
| Sex (female) | 26,017 | 0.564 | 0.496 | 0 | 1 | 10,028 | 0.531 | 0.499 | 0 | 1 |

### 4 Equations and regression tables corresponding to main results

This section shows regression tables that correspond to the main results figures, using both a birth period dummy approach, and a continuous birth year approach. The two types of models are estimated between families and within families. Before showing the tables, the model equations are shown.

#### 4.1 Equations

*Interaction between EA PGI and birth cohort*

Beginning with the interaction between the EA PGI and birth cohort, using the main sample (twin cohorts born 1925–1989), the following model is estimated *between families*:

$$Edu_{ij} = \alpha + \beta_1(EAPGI_{ij} \times BY_{ij}) + \beta_2 EAPGI_{ij} + \beta_3 BY_{ij} + \Lambda_{ij} + \epsilon_{ij} \tag{2}$$

where $Edu^c_{ij}$ represents education decile for twin $i$ in twin pair $j$. The interaction between the EA PGI and birth cohort is captured by the term $EAPGI_{ij} \times BY_{ij}$, where $BY^c_{ij}$ can take the categorical values 0 (1925–1949), 1 (1950–1969), or 3 (1970–1989), or, in the continuous birth year approach, 1925–1989. $\Lambda$ includes the covariates (the first 20 principal components of the genetic data, and interactions between birth year and sex) interacted with the main independent variables as well as with each other. Standard errors are clustered on the level of twin pair.

*Within families*, the following model is estimated:

$$\Delta Edu^c_j = \gamma + \theta_1(\Delta EAPGI_j \times BY_j) + \theta_2 \Delta EAPGI_j + \theta_3 BY_j + \Omega_j + \epsilon_j \tag{3}$$

where the object of analysis goes from being twin $i$ to being twin pair $j$, and where $\Delta Edu^c_j$ represents the *difference* in education decile between two dizygotic twins in a dizygotic twin pair. The interaction between the EA PGI and birth cohort is captured by $\Delta EAPGI_j \times BY_j$, where $\Delta EAPGI_j$ represents the twin difference in the EA PGI, and where $BY_j$, as in (1), can take the categorical values 0 (1925–1949), 1 (1950–1969), or 3 (1970–1989), or the continuous values 1925–1989. The $\Omega_j$ term includes a covariate for sex difference within twin pair, and interactions between the covariate and the main independent variables.

*Interaction between EA PGI, parental education, and birth cohort*

Continuing to the three-way interaction between the EA PGI, parental education, and birth cohort (i.e. testing whether the interaction between the EA PGI and parental education changes over time), now using the parental sample (twin cohorts born 1940–1989), the following model is estimated *between families*:

$$\begin{aligned} Edu^c_{ij} = \alpha &+ \beta_1(EAPGI^c_{ij} \times Edu^p_{ij} \times BY^c_{ij}) + \beta_2(EAPGI^c_{ij} \times Edu^p_{ij}) \\ &+\beta_3(EAPGI^c_{ij} \times BY^c_{ij}) + \beta_4(Edu^p_{ij} \times BY^c_{ij}) \\ &+\beta_5(EAPGI_{ij}) + \beta_6(Edu^p_{ij}) + \beta_7(BY^c_{ij}) + \Lambda_{ij} + \epsilon_{ij} \end{aligned} \tag{4}$$

where the three-way interaction is captured by the term $EAPGI_{ij}^{c} \times Edu_{ij}^{p} \times BY_{ij}^{c}$. Superscripts $^{c}$ (for child) and $^{p}$ (for parent) are included to distinguish twin variables from parental variables. Written out are also each possible two-way interaction between the main independent variables, and the constitutive terms of each interaction, all of which need to be included in the model. The $\Lambda_{ij}$ term includes the same covariates as in (2), plus all possible interactions between the covariates and the main independent variables, and with eachother.

*Within families*, the following model is estimated:

$$
\begin{aligned}
\Delta Edu_{j}^{c} = \alpha + \beta_1(\Delta EAPGI_{j}^{c} \times Edu_{j}^{p} \times BY_{j}^{c}) + \theta_2(\Delta EAPGI_{j}^{c} \times Edu_{j}^{p}) \\
+\theta_3(\Delta EAPGI_{j}^{c} \times BY_{j}^{c}) + \theta_4(Edu_{j}^{p} \times BY_{j}^{c}) \\
+\theta_5(\Delta EAPGI_{j}) + \theta_6(Edu_{j}^{p}) + \theta_7(BY_{j}^{c}) + \Omega_j + \epsilon
\end{aligned}
\tag{5}
$$

where the three-way interaction is captured by the term $\Delta EAPGI_{j}^{c} \times Edu_{j}^{p} \times BY_{j}^{c}$. The $\Omega_j$ term includes all possible interactions between covariates (sex difference) and the main independent variables.

#### 4.2 Regression tables for multi-trait EA PGI analyses

Table A4: Interaction between EA PGI and birth year for educational attainment, 1925–1989 (dummies)

a) Between family

| VARIABLES | (1) | (2) | (3) |
|---|---|---|---|
| EA PGI (multi) | 0.699*** | | 0.561*** |
| | (0.014) | | (0.026) |
| 1950-1969 | | -0.299*** | -0.274*** |
| | | (0.037) | (0.050) |
| 1970-1989 | | 0.401*** | 0.479*** |
| | | (0.039) | (0.052) |
| EA PGI (multi) x 1950-1969 | | | 0.112*** |
| | | | (0.034) |
| EA PGI (multi) x 1970-1989 | | | 0.161*** |
| | | | (0.035) |
| Constant | 7.094*** | 7.141*** | 7.095*** |
| | (0.032) | (0.029) | (0.032) |
| Observations | 28,628 | 29,417 | 28,628 |
| R-squared | 0.105 | 0.015 | 0.110 |

b) Within family

| VARIABLES | (1) | (2) | (3) |
|---|---|---|---|
| Δ EA PGI (multi) | 0.383*** | | 0.323*** |
| | (0.037) | | (0.050) |
| 1950-1969 | | 0.015 | 0.020 |
| | | (0.077) | (0.078) |
| 1970-1989 | | -0.001 | 0.010 |
| | | (0.090) | (0.091) |
| Δ EA PGI (multi) x 1950-1969 | | | 0.124 |
| | | | (0.084) |
| Δ EA PGI (multi) x 1970-1989 | | | 0.141 |
| | | | (0.102) |
| Constant | 0.013 | 0.004 | 0.005 |
| | (0.034) | (0.047) | (0.047) |
| Observations | 5,415 | 5,565 | 5,415 |
| R-squared | 0.021 | 0.001 | 0.021 |

Note: Each between-family model includes controls for the first 20 principal components of the genetic data, and an interaction between birth year and sex. Each within-family model includes a control for sex difference within a twin pair. In the interaction models, all controls are interacted with the independent variables. Standard errors, shown in parentheses, allow for clustering at twin-pair level. *** p<0.01, ** p<0.05, * p<0.1

Table A5: Interaction between EA PGI and birth year for educational attainment, 1925–1989 (continuous)

a) Between family

| VARIABLES | (1) | (2) | (3) |
|---|---|---|---|
| EA PGI (multi) | 0.699*** | | -7.676*** |
| | (0.014) | | (1.611) |
| Birth year | | 0.007*** | 0.008*** |
| | | (0.001) | (0.001) |
| EA PGI (multi) x Birth year | | | 0.004*** |
| | | | (0.001) |
| Constant | -8.049*** | -6.120*** | -7.960*** |
| | (2.284) | (1.761) | (2.284) |
| Observations | 28,628 | 29,417 | 28,628 |
| R-squared | 0.094 | 0.004 | 0.098 |

b) Within family

| VARIABLES | (1) | (2) | (3) |
|---|---|---|---|
| Δ EA PGI (multi) | 0.383*** | | -9.053** |
| | (0.037) | | (4.394) |
| Birth year | | 0.001 | |
| | | (0.002) | |
| Δ EA PGI (multi) x Birth year | | | 0.005** |
| | | | (0.002) |
| Constant | 0.013 | -1.748 | -0.217 |
| | (0.034) | (3.918) | (0.452) |
| Observations | 5,415 | 5,565 | 5,415 |
| R-squared | 0.021 | 0.001 | 0.051 |

Note: Each between-family model includes controls for the first 20 principal components of the genetic data, and an interaction between birth year and sex. Each within-family model includes a control for sex difference within a twin pair. In the interaction models, all controls are interacted with the independent variables. Standard errors, shown in parentheses, allow for clustering at twin-pair level. *** $p<0.01$, ** $p<0.05$, * $p<0.1$

Table A6: Interaction between EA PGI, parental education, and birth year for educational attainment, 1940–1989 (dummies)

a) Between family

| VARIABLES | (1) | (2) | (3) | (4) | (5) |
|---|---|---|---|---|---|
| EA PGI (multi) | 0.732*** | | 0.275*** | | -0.351 |
| | (0.016) | | (0.094) | | (0.316) |
| Parent edu. decile | | 0.415*** | 0.539*** | | 0.544*** |
| | | (0.010) | (0.039) | | (0.039) |
| 1950-1969 | | | | -0.058 | 2.039*** |
| | | | | (0.043) | (0.364) |
| 1970-1989 | | | | 0.642*** | 3.124*** |
| | | | | (0.044) | (0.355) |
| EA PGI (multi) x 1950-1969 | | | | | 0.564* |
| | | | | | (0.340) |
| EA PGI (multi) x 1970-1989 | | | | | 0.790** |
| | | | | | (0.333) |
| Parent edu. decile x 1950-1969 | | | | | -0.214*** |
| | | | | | (0.041) |
| Parent edu. decile x 1970-1989 | | | | | -0.245*** |
| | | | | | (0.040) |
| EA PGI (multi) x Parent edu. decile | | | 0.033*** | | 0.103*** |
| | | | (0.010) | | (0.035) |
| EA PGI (multi) x Parent edu. decile x 1950-1969 | | | | | -0.060 |
| | | | | | (0.038) |
| EA PGI (multi) x Parent edu. decile x 1970-1989 | | | | | -0.091** |
| | | | | | (0.037) |
| Constant | -37.370*** | 3.128*** | 2.088*** | 6.893*** | 2.039*** |
| | (2.978) | (0.104) | (0.346) | (0.037) | (0.346) |
| Observations | 24,943 | 19,970 | 19,750 | 25,674 | 19,750 |
| R-squared | 0.110 | 0.103 | 0.175 | 0.019 | 0.175 |

b) Within family

| VARIABLES | (1) | (2) | (3) | (4) | (5) |
|---|---|---|---|---|---|
| Δ EA PGI (multi) | 0.436*** | | 0.630** | | -1.849** |
| | (0.043) | | (0.252) | | (0.858) |
| Parent edu. decile | | 0.013 | 0.009 | | 0.001 |
| | | (0.028) | (0.028) | | (0.085) |
| 1950-1969 | | | | 0.062 | -0.024 |
| | | | | (0.091) | (0.841) |
| 1970-1989 | | | | 0.045 | -0.049 |
| | | | | (0.103) | (0.825) |
| Δ EA PGI (multi) x 1950-1969 | | | | | 2.704*** |
| | | | | | (0.935) |
| Δ EA PGI (multi) x 1970-1989 | | | | | 2.711*** |
| | | | | | (0.937) |
| Parent edu. decile x 1950-1969 | | | | | 0.011 |
| | | | | | (0.096) |
| Parent edu. decile x 1970-1989 | | | | | 0.013 |
| | | | | | (0.094) |
| Δ EA PGI (multi) x Parent edu. decile | | | -0.025 | | 0.252*** |
| | | | (0.030) | | (0.097) |
| Δ EA PGI (multi) x Parent edu. decile x 1950-1969 | | | | | -0.303*** |
| | | | | | (0.107) |
| Δ EA PGI (multi) x Parent edu. decile x 1970-1989 | | | | | -0.302*** |
| | | | | | (0.107) |
| Constant | 0.004 | -0.091 | -0.051 | -0.043 | -0.033 |
| | (0.040) | (0.234) | (0.235) | (0.065) | (0.754) |
| Observations | 4,264 | 3,711 | 3,636 | 4,408 | 3,636 |
| R-squared | 0.024 | 0.001 | 0.023 | 0.001 | 0.026 |

Note: Each between-family model includes controls for the first 20 principal components of the genetic data, and an interaction between birth year and sex. Each within-family model includes a control for sex difference within a twin pair. In the interaction models, all controls are interacted with the independent variables. Standard errors, shown in parentheses, allow for clustering at twin-pair level. *** p<0.01, ** p<0.05, * p<0.1

Table A7: Interaction between EA PGI, parental education, and birth year for educational attainment, 1940–1989 (continuous)

a) Between family

| VARIABLES | (1) | (2) | (3) | (4) | (5) |
|---|---|---|---|---|---|
| EA PGI (multi) | 0.732*** | | 1.297 | | -25.304** |
| | (0.016) | | (2.294) | | (12.115) |
| Parent edu. decile | | 0.426*** | 10.202*** | | 10.600*** |
| | | (0.010) | (1.492) | | (1.494) |
| Birth year | | | | 0.019*** | 0.074*** |
| | | | | (0.001) | (0.007) |
| EA PGI (multi) x Parent edu. decile | | | 0.029*** | | 3.183** |
| | | | (0.010) | | (1.383) |
| EA PGI (multi) x Parent edu. decile x Birth year | | | | | -0.002** |
| | | | | | (0.001) |
| Constant | -37.370*** | -69.158*** | -136.509*** | -30.400*** | -140.864*** |
| | (2.978) | (3.307) | (12.816) | (2.221) | (12.901) |
| Observations | 24,943 | 19,970 | 19,750 | 25,674 | 19,750 |
| R-squared | 0.110 | 0.105 | 0.174 | 0.016 | 0.174 |

b) Within family

| VARIABLES | (1) | (2) | (3) | (4) | (5) |
|---|---|---|---|---|---|
| Δ EA PGI (multi) | 0.436*** | | 0.630** | | -46.445 |
| | (0.043) | | (0.252) | | (35.374) |
| Parent edu. decile | | 0.013 | 0.009 | | 2.583 |
| | | (0.028) | (0.028) | | (3.787) |
| Birth year | | | | 0.003 | 0.015 |
| | | | | (0.003) | (0.016) |
| Δ EA PGI (multi) x Parent edu. decile | | | -0.025 | | 4.934 |
| | | | (0.030) | | (4.147) |
| Δ EA PGI (multi) x Parent edu. decile x Birth year | | | | | -0.003 |
| | | | | | (0.002) |
| Constant | 0.004 | -0.091 | -0.051 | -6.202 | -30.263 |
| | (0.040) | (0.234) | (0.235) | (5.212) | (32.392) |
| Observations | 4,264 | 3,711 | 3,636 | 4,408 | 3,636 |
| R-squared | 0.024 | 0.001 | 0.023 | 0.001 | 0.024 |

Note: Each between-family model includes controls for the first 20 principal components of the genetic data, and an interaction between birth year and sex. Each within-family model includes a control for sex difference within a twin pair. In the interaction models, all controls are interacted with the independent variables. Standard errors, shown in parentheses, allow for clustering at twin-pair level. *** p<0.01, ** p<0.05, * p<0.1

### 5 Robustness tests for main analysis

#### 5.1 Coefficient plots and regression tables for single-trait EA PGI analyses

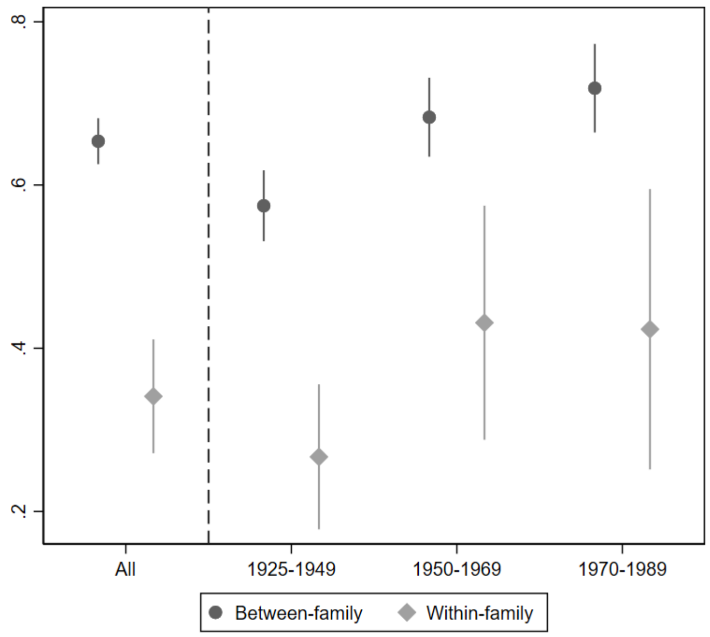

Figure A2: Relationship between EA PGI and educational attainment by birth cohort

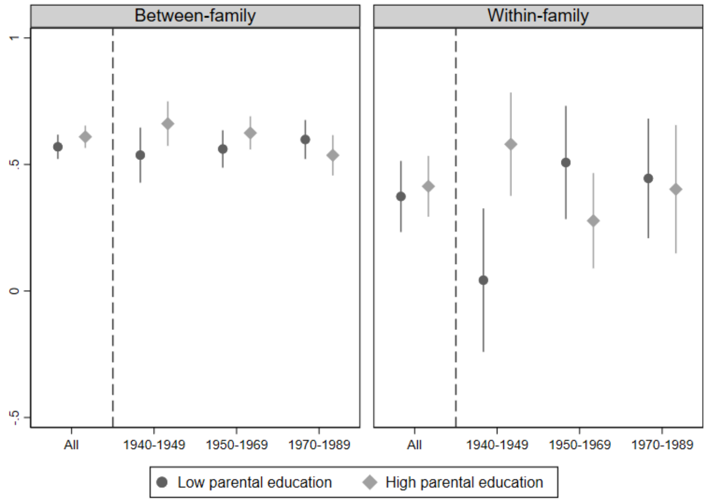

Figure A3: Relationship between EA PGI and educational attainment by birth cohort, split by parental education

Table A8: Interaction between EA PGI and birth year for educational attainment, 1925–1989 (dummies)

a) Between family

| VARIABLES | (1) | (2) | (3) |
|---|---|---|---|
| EA PGI (single) | 0.654*** | | 0.527*** |
| | (0.014) | | (0.026) |
| 1950-1969 | | -0.299*** | -0.273*** |
| | | (0.037) | (0.051) |
| 1970-1989 | | 0.401*** | 0.482*** |
| | | (0.039) | (0.052) |
| EA PGI (single) x 1950-1969 | | | 0.102*** |
| | | | (0.034) |
| EA PGI (single) x 1970-1989 | | | 0.139*** |
| | | | (0.036) |
| Constant | 7.093*** | 7.141*** | 7.095*** |
| | (0.032) | (0.029) | (0.032) |
| Observations | 28,628 | 29,417 | 28,628 |
| R-squared | 0.094 | 0.015 | 0.099 |

b) Within family

| VARIABLES | (1) | (2) | (3) |
|---|---|---|---|
| Δ EA PGI (single) | 0.341*** | | 0.267*** |
| | (0.036) | | (0.049) |
| 1950-1969 | | 0.015 | 0.024 |
| | | (0.077) | (0.078) |
| 1970-1989 | | -0.001 | 0.006 |
| | | (0.090) | (0.091) |
| Δ EA PGI (single) x 1950-1969 | | | 0.156* |
| | | | (0.083) |
| Δ EA PGI (single) x 1970-1989 | | | 0.159 |
| | | | (0.099) |
| Constant | 0.013 | 0.004 | 0.005 |
| | (0.034) | (0.047) | (0.047) |
| Observations | 5,415 | 5,565 | 5,415 |
| R-squared | 0.017 | 0.001 | 0.018 |

Note: Each between-family model includes controls for the first 20 principal components of the genetic data, and an interaction between birth year and sex. Each within-family model includes a control for sex difference within a twin pair. In the interaction models, all controls are interacted with the independent variables. Standard errors, shown in parentheses, allow for clustering at twin-pair level. *** p<0.01, ** p<0.05, * p<0.1

Table A9: Interaction between EA PGI and birth year for educational attainment, 1925–1989 (continuous)

a) Between family

| VARIABLES | (1) | (2) | (3) |
|---|---|---|---|
| EA PGI (single) | 0.654*** | | -6.397*** |
| | (0.014) | | (1.631) |
| Birth year | | 0.007*** | 0.008*** |
| | | (0.001) | (0.001) |
| EA PGI (single) x Birth year | | | 0.004*** |
| | | | (0.001) |
| Constant | -8.278*** | -6.120*** | -8.147*** |
| | (2.298) | (1.761) | (2.298) |
| Observations | 28,628 | 29,417 | 28,628 |
| R-squared | 0.083 | 0.004 | 0.087 |

b) Within family

| VARIABLES | (1) | (2) | (3) |
|---|---|---|---|
| Δ EA PGI (single) | 0.341*** | | -10.094** |
| | (0.036) | | (4.279) |
| Birth year | | 0.001 | |
| | | (0.002) | |
| Δ EA PGI (single) x Birth year | | | 0.005** |
| | | | (0.002) |
| Constant | 0.013 | -1.748 | -0.178 |
| | (0.034) | (3.918) | (0.453) |
| Observations | 5,415 | 5,565 | 5,415 |
| R-squared | 0.017 | 0.001 | 0.048 |

Note: Each between-family model includes controls for the first 20 principal components of the genetic data, and an interaction between birth year and sex. Each within-family model includes a control for sex difference within a twin pair. In the interaction models, all controls are interacted with the independent variables. Standard errors, shown in parentheses, allow for clustering at twin-pair level. *** $p<0.01$, ** $p<0.05$, * $p<0.1$

Table A10: Interaction between EA PGI, parental education, and birth year for educational attainment, 1940–1989 (dummies)

a) Between family

| VARIABLES | (1) | (2) | (3) | (4) | (5) |
|---|---|---|---|---|---|
| EA PGI (single) | 0.683*** | | 0.257*** | | -0.460 |
| | (0.016) | | (0.092) | | (0.307) |
| Parent edu. decile | | 0.415*** | 0.547*** | | 0.552*** |
| | | (0.010) | (0.039) | | (0.039) |
| 1950-1969 | | | | -0.058 | 2.060*** |
| | | | | (0.043) | (0.365) |
| 1970-1989 | | | | 0.642*** | 3.126*** |
| | | | | (0.044) | (0.356) |
| EA PGI (single) x 1950-1969 | | | | | 0.684** |
| | | | | | (0.332) |
| EA PGI (single) x 1970-1989 | | | | | 0.853*** |
| | | | | | (0.324) |
| Parent edu. decile x 1950-1969 | | | | | -0.215*** |
| | | | | | (0.041) |
| Parent edu. decile x 1970-1989 | | | | | -0.244*** |
| | | | | | (0.040) |
| EA PGI (single) x Parent edu. decile | | | 0.031*** | | 0.113*** |
| | | | (0.009) | | (0.034) |
| EA PGI (single) x Parent edu. decile x 1950-1969 | | | | | -0.075** |
| | | | | | (0.037) |
| EA PGI (single) x Parent edu. decile x 1970-1989 | | | | | -0.100*** |
| | | | | | (0.036) |
| Constant | -37.361*** | 3.128*** | 2.022*** | 6.893*** | 1.965*** |
| | (2.998) | (0.104) | (0.346) | (0.037) | (0.346) |
| Observations | 24,943 | 19,970 | 19,750 | 25,674 | 19,750 |
| R-squared | 0.098 | 0.103 | 0.166 | 0.019 | 0.166 |

b) Within family

| VARIABLES | (1) | (2) | (3) | (4) | (5) |
|---|---|---|---|---|---|
| Parent edu. decile | | 0.013 | 0.004 | | -0.005 |
| | | (0.028) | (0.028) | | (0.085) |
| Δ EA PGI (single) | 0.398*** | | 0.479** | | -2.114** |
| | (0.043) | | (0.244) | | (0.835) |
| 1950-1969 | | | | 0.062 | -0.005 |
| | | | | (0.091) | (0.842) |
| 1970-1989 | | | | 0.045 | -0.044 |
| | | | | (0.103) | (0.826) |
| Δ EA PGI (single) x 1950-1969 | | | | | 3.028*** |
| | | | | | (0.912) |
| Δ EA PGI (single) x 1970-1989 | | | | | 2.641*** |
| | | | | | (0.909) |
| Parent edu. decile x 1950-1969 | | | | | 0.010 |
| | | | | | (0.096) |
| Parent edu. decile x 1970-1989 | | | | | 0.012 |
| | | | | | (0.094) |
| Δ EA PGI (single) x Parent edu. decile | | | -0.010 | | 0.279*** |
| | | | (0.029) | | (0.094) |
| Δ EA PGI (single) x Parent edu. decile x 1950-1969 | | | | | -0.341*** |
| | | | | | (0.104) |
| Δ EA PGI (single) x Parent edu. decile x 1970-1989 | | | | | -0.292*** |
| | | | | | (0.104) |
| Constant | 0.002 | -0.091 | -0.014 | -0.043 | 0.011 |
| | (0.040) | (0.234) | (0.235) | (0.065) | (0.755) |
| Observations | 4,264 | 3,711 | 3,636 | 4,408 | 3,636 |
| R-squared | 0.021 | 0.001 | 0.021 | 0.001 | 0.024 |

Note: Each between-family model includes controls for the first 20 principal components of the genetic data, and an interaction between birth year and sex. Each within-family model includes a control for sex difference within a twin pair. In the interaction models, all controls are interacted with the independent variables. Standard errors, shown in parentheses, allow for clustering at twin-pair level. *** p<0.01, ** p<0.05, * p<0.1

Table A11: Interaction between EA PGI, parental education, and birth year for educational attainment, 1940–1989 (continuous)

a) Between family

| VARIABLES | (1) | (2) | (3) | (4) | (5) |
|---|---|---|---|---|---|
| EA PGI (single) | 0.683*** | | 2.184 | | -25.586** |
| | (0.016) | | (2.307) | | (11.903) |
| Parent edu. decile | | 0.426*** | 9.976*** | | 10.342*** |
| | | (0.010) | (1.494) | | (1.498) |
| Birth year | | | | 0.019*** | 0.073*** |
| | | | | (0.001) | (0.007) |
| EA PGI (single) x Parent edu. decile | | | 0.028*** | | 3.339** |
| | | | (0.009) | | (1.366) |
| EA PGI (single) x Parent edu. decile x Birth year | | | | | -0.002** |
| | | | | | (0.001) |
| Constant | -37.361*** | -69.158*** | -135.274*** | -30.400*** | -139.343*** |
| | (2.998) | (3.307) | (12.844) | (2.221) | (12.938) |
| Observations | 24,943 | 19,970 | 19,750 | 25,674 | 19,750 |
| R-squared | 0.098 | 0.105 | 0.166 | 0.016 | 0.166 |

b) Within family

| VARIABLES | (1) | (2) | (3) | (4) | (5) |
|---|---|---|---|---|---|
| Parent edu. decile | | 0.013 | 0.004 | | 2.737 |
| | | (0.028) | (0.028) | | (3.794) |
| Δ EA PGI (single) | 0.398*** | | 0.479** | | -20.665 |
| | (0.043) | | (0.244) | | (33.793) |
| Birth year | | | | 0.003 | 0.016 |
| | | | | (0.003) | (0.016) |
| Δ EA PGI (single) x Parent edu. decile | | | -0.010 | | 1.887 |
| | | | (0.029) | | (3.965) |
| Δ EA PGI (single) x Parent edu. decile x Birth year | | | | | -0.001 |
| | | | | | (0.002) |
| Constant | 0.002 | -0.091 | -0.014 | -6.202 | -31.129 |
| | (0.040) | (0.234) | (0.235) | (5.212) | (32.448) |
| Observations | 4,264 | 3,711 | 3,636 | 4,408 | 3,636 |
| R-squared | 0.021 | 0.001 | 0.021 | 0.001 | 0.022 |

Note: Each between-family model includes controls for the first 20 principal components of the genetic data, and an interaction between birth year and sex. Each within-family model includes a control for sex difference within a twin pair. In the interaction models, all controls are interacted with the independent variables. Standard errors, shown in parentheses, allow for clustering at twin-pair level. *** p<0.01, ** p<0.05, * p<0.1

#### 5.2 Alternative operationalisations of parental education

This section shows results corresponding to figure 3 in the main text, using alternative specifications of parental education. Specifically, the models are run using either the mother's highest education, or the father's highest education. Corresponding regression tables are available on request.

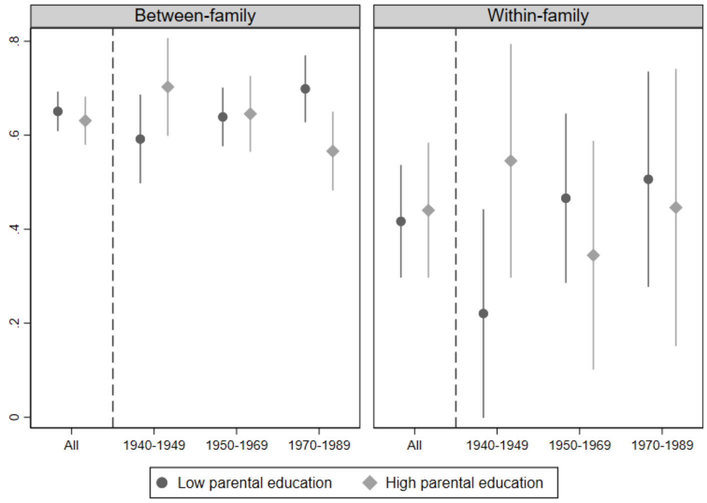

Figure A4: Effect of EA PGI by birth cohort, split on mother's education

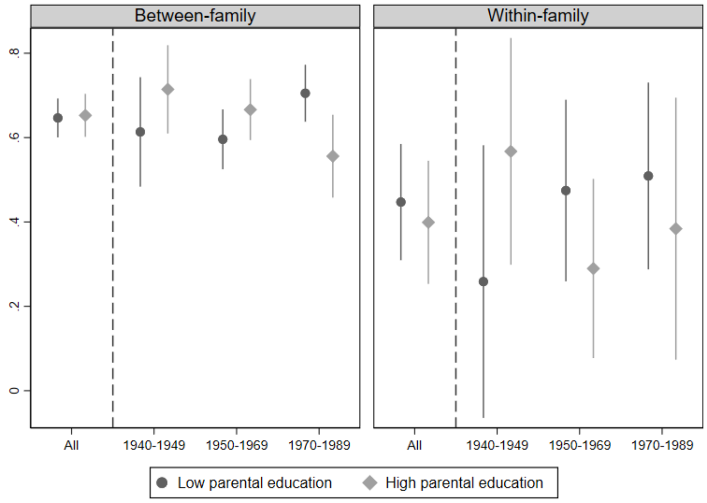

Figure A5: Effect of EA PGI by birth cohort, split on father's education

#### 5.3 Comparing PGI models using only complete DZ twin pairs

This section shows graphs corresponding to figures 2-3 in the main text, comparing the between-family model with the within-family model, but restricts the comparison to full DZ twin pairs only. Since the within-family model by definition requires complete twin pairs, the estimates for this model will be the same. The sample changes for the between-family model, however, since the sample can also encompass MZ twins and singleton DZ twins. Corresponding regression tables are available on request.

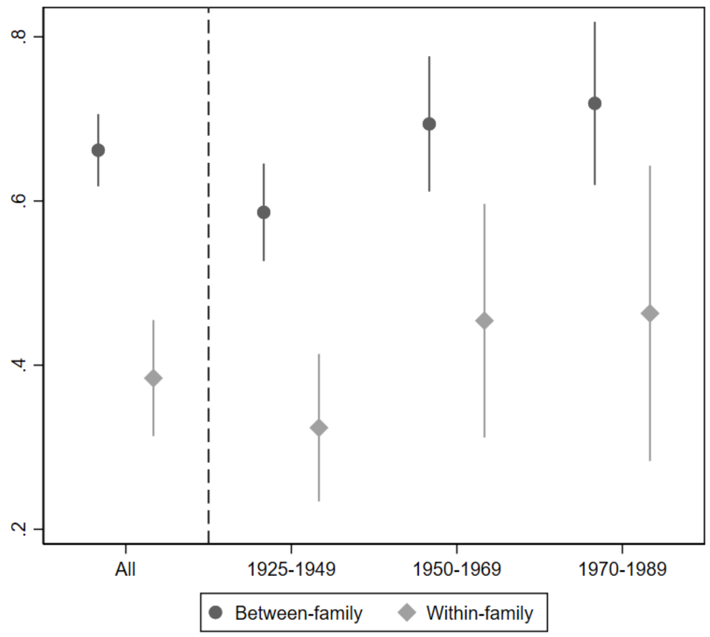

Figure A6: Effect of EA PGI by birth cohort (Complete DZ pairs only)

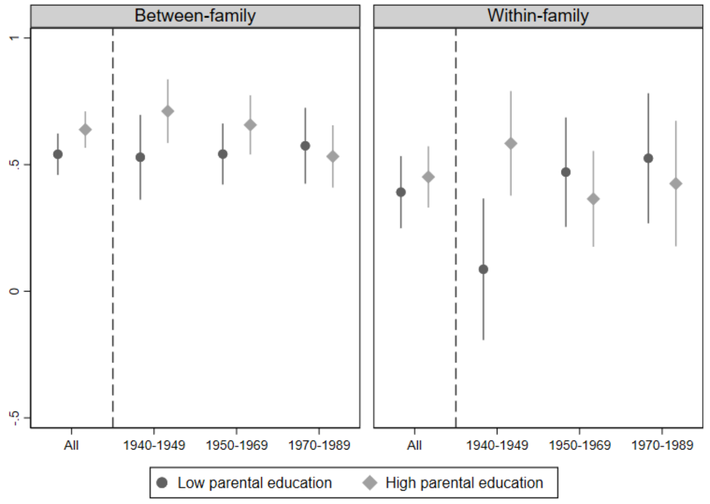

Figure A7: Effect of EA PGI by birth cohort, split on parental education (Full DZ pairs only)

### 6 Results for supplementary dependent variables

#### 6.1 Tertiary education

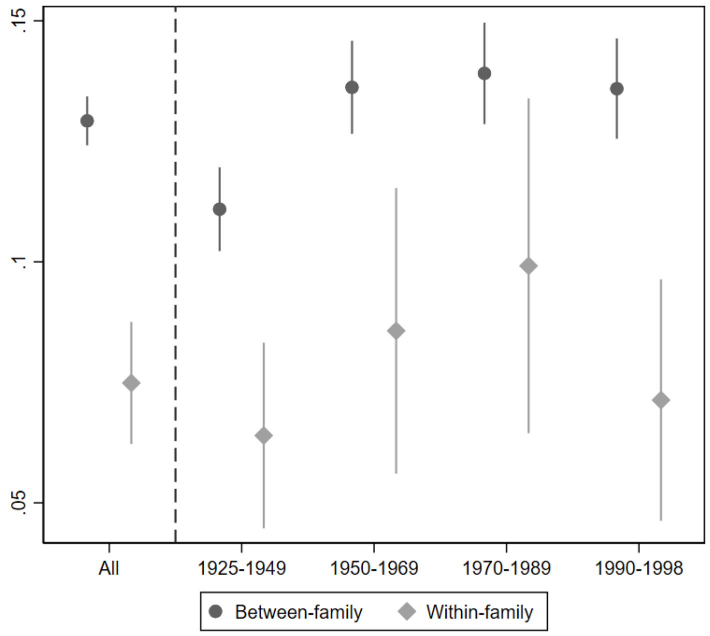

Figure A8: Effect of EA PGI on having tertiary education by birth cohort

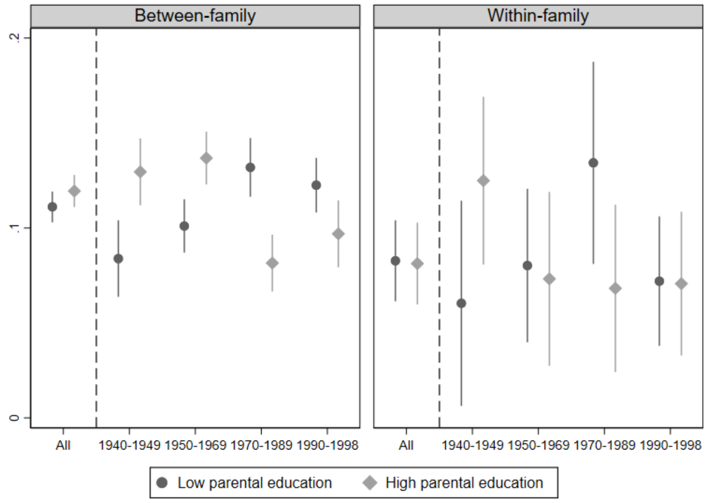

Figure A9: Effect of EA PGI on having tertiary education by birth cohort, split on parental education

Table A12: Interaction between EA PGI and birth year for tertiary education, 1925–1998 (dummies)

a) Between family

| VARIABLES | (1) | (2) | (3) |
|---|---|---|---|
| EA PGI (multi) | 0.129*** | | 0.112*** |
| | (0.003) | | (0.005) |
| 1950-1969 | | 0.150*** | 0.132*** |
| | | (0.007) | (0.010) |
| 1970-1989 | | 0.392*** | 0.347*** |
| | | (0.008) | (0.011) |
| 1990-1998 | | 0.146*** | 0.084*** |
| | | (0.008) | (0.010) |
| EA PGI (multi) x 1950-1969 | | | 0.025*** |
| | | | (0.007) |
| EA PGI (multi) x 1970-1989 | | | 0.029*** |
| | | | (0.007) |
| EA PGI (multi) x 1990-1998 | | | 0.024*** |
| | | | (0.007) |
| Constant | 0.259*** | 0.235*** | 0.260*** |
| | (0.006) | (0.006) | (0.006) |
| Observations | 36,912 | 38,080 | 36,912 |
| R-squared | 0.156 | 0.086 | 0.161 |

b) Within family

| VARIABLES | (1) | (2) | (3) |
|---|---|---|---|
| Δ EA PGI (multi) | 0.075*** | | 0.065*** |
| | (0.007) | | (0.011) |
| 1950-1969 | | 0.020 | 0.016 |
| | | (0.016) | (0.016) |
| 1970-1989 | | 0.001 | -0.001 |
| | | (0.019) | (0.019) |
| 1990-1998 | | -0.009 | -0.010 |
| | | (0.015) | (0.015) |
| Δ EA PGI (multi) x 1950-1969 | | | 0.018 |
| | | | (0.018) |
| Δ EA PGI (multi) x 1970-1989 | | | 0.033 |
| | | | (0.021) |
| Δ EA PGI (multi) x 1990-1989 | | | 0.006 |
| | | | (0.016) |
| Constant | -0.002 | -0.004 | -0.002 |
| | (0.006) | (0.010) | (0.010) |
| Observations | 7,492 | 7,764 | 7,492 |
| R-squared | 0.023 | 0.006 | 0.028 |

Note: Each between-family model includes controls for the first 20 principal components of the genetic data, and an interaction between birth year and sex. Each within-family model includes a control for sex difference within a twin pair. In the interaction models, all controls are interacted with the independent variables. Standard errors, shown in parentheses, allow for clustering at twin-pair level. *** $p<0.01$, ** $p<0.05$, * $p<0.1$

Table A13: Interaction between EA PGI and birth year for tertiary education, 1925–1998 (linear)

a) Between family

| VARIABLES | (1) | (2) | (3) |
| --- | --- | --- | --- |
| EA PGI (multi) | 0.129*** | | -0.679*** |
| | (0.003) | | (0.241) |
| Birth year | | 0.004*** | 0.003*** |
| | | (0.000) | (0.000) |
| EA PGI (multi) x Birth year | | | 0.000*** |
| | | | (0.000) |
| Constant | -5.357*** | -7.837*** | -5.314*** |
| | (0.354) | (0.266) | (0.354) |
| Observations | 36,912 | 38,080 | 36,912 |
| R-squared | 0.112 | 0.040 | 0.114 |

b) Within family

| VARIABLES | (1) | (2) | (3) |
| --- | --- | --- | --- |
| Δ EA PGI (multi) | 0.075*** | | -1.727* |
| | (0.007) | | (0.911) |
| Birth year | | -0.000 | 0.001 |
| | | (0.000) | (0.001) |
| Δ EA PGI (multi) x Birth year | | | 0.001** |
| | | | (0.000) |
| Constant | -0.002 | 0.341 | -1.494 |
| | (0.006) | (0.502) | (2.246) |
| Observations | 7,492 | 7,764 | 5,412 |
| R-squared | 0.023 | 0.006 | 0.027 |

Note: Each between-family model includes controls for the first 20 principal components of the genetic data, and an interaction between birth year and sex. Each within-family model includes a control for sex difference within a twin pair. In the interaction models, all controls are interacted with the independent variables. Standard errors, shown in parentheses, allow for clustering at twin-pair level. *** p<0.01, ** p<0.05, * p<0.1

Table A14: Interaction between EA PGI and birth year for tertiary education, 1940–1998 (dummies)

a) Between family

| VARIABLES | (1) | (2) | (3) | (4) | (5) |
|---|---|---|---|---|---|
| EA PGI (multi) | 0.133*** | | 0.068*** | | -0.171*** |
| | (0.003) | | (0.015) | | (0.061) |
| Parent edu. decile | | 0.070*** | 0.094*** | | 0.095*** |
| | | (0.002) | (0.008) | | (0.008) |
| 1950-1969 | | | | 0.130*** | 0.409*** |
| | | | | (0.008) | (0.071) |
| 1970-1989 | | | | 0.371*** | 0.684*** |
| | | | | (0.008) | (0.069) |
| 1990-1998 | | | | 0.127*** | 0.566*** |
| | | | | (0.009) | (0.069) |
| EA PGI (multi) x 1950-1969 | | | | | 0.173*** |
| | | | | | (0.066) |
| EA PGI (multi) x 1970-1989 | | | | | 0.316*** |
| | | | | | (0.065) |
| EA PGI (multi) x 1990-1998 | | | | | 0.252*** |
| | | | | | (0.065) |
| Parent edu. decile x 1950-1969 | | | | | -0.029*** |
| | | | | | (0.008) |
| Parent edu. decile x 1970-1989 | | | | | -0.029*** |
| | | | | | (0.008) |
| Parent edu. decile x 1990-1998 | | | | | -0.045*** |
| | | | | | (0.008) |
| EA PGI (multi) x Parent edu. decile | | | 0.005*** | | 0.032*** |
| | | | (0.002) | | (0.007) |
| EA PGI (multi) x Parent edu. decile x 1950-1969 | | | | | -0.018** |
| | | | | | (0.008) |
| EA PGI (multi) x Parent edu. decile x 1970-1989 | | | | | -0.037*** |
| | | | | | (0.007) |
| EA PGI (multi) x Parent edu. decile x 1990-1998 | | | | | -0.029*** |
| | | | | | (0.007) |
| Constant | -4.068*** | -0.327*** | -0.564*** | 0.251*** | -0.583*** |
| | (0.426) | (0.018) | (0.065) | (0.007) | (0.066) |
| Observations | 33,231 | 26,639 | 26,298 | 34,341 | 26,298 |
| R-squared | 0.101 | 0.135 | 0.187 | 0.076 | 0.188 |

b) Within family

| VARIABLES | (1) | (2) | (3) | (4) | (5) |
|---|---|---|---|---|---|
| Δ EA PGI (multi) | 0.082*** | | 0.074** | | -0.291* |
| | (0.007) | | (0.036) | | (0.176) |
| Parent edu. decile | | -0.001 | -0.002 | | -0.026 |
| | | (0.004) | (0.004) | | (0.018) |
| 1950-1969 | | | | 0.029 | -0.140 |
| | | | | (0.019) | (0.173) |
| 1970-1989 | | | | 0.009 | -0.255 |
| | | | | (0.021) | (0.170) |
| 1990-1998 | | | | -0.000 | -0.232 |
| | | | | (0.018) | (0.162) |
| Δ EA PGI (multi) x 1950-1969 | | | | | 0.421** |
| | | | | | (0.192) |
| Δ EA PGI (multi) x 1970-1989 | | | | | 0.470** |
| | | | | | (0.193) |
| Δ EA PGI (multi) x 1990-1998 | | | | | 0.334* |
| | | | | | (0.184) |
| Parent edu. decile x 1950-1969 | | | | | 0.019 |
| | | | | | (0.020) |
| Parent edu. decile x 1970-1989 | | | | | 0.030 |
| | | | | | (0.019) |
| Parent edu. decile x 1990-1998 | | | | | 0.026 |
| | | | | | (0.018) |
| Δ EA PGI (multi) x Parent edu. decile | | | 0.001 | | 0.044** |
| | | | (0.004) | | (0.020) |
| Δ EA PGI (multi) x Parent edu. decile x 1950-1969 | | | | | -0.050** |
| | | | | | (0.022) |
| Δ EA PGI (multi) x Parent edu. decile x 1970-1989 | | | | | -0.054** |
| | | | | | (0.022) |
| Δ EA PGI (multi) x Parent edu. decile x 1990-1998 | | | | | -0.040* |
| | | | | | (0.021) |
| Constant | -0.002 | 0.009 | 0.011 | -0.013 | 0.220 |
| | (0.007) | (0.033) | (0.033) | (0.013) | (0.155) |
| Observations | 6,343 | 5,824 | 5,684 | 6,609 | 5,684 |
| R-squared | 0.028 | 0.008 | 0.028 | 0.009 | 0.035 |

Note: Each between-family model includes controls for the first 20 principal components of the genetic data, and an interaction between birth year and sex. Each within-family model includes a control for sex difference within a twin pair. In the [illegible]

Table A15: Interaction between EA PGI and birth year for tertiary education, 1940–1998 (dummies)

a) Between family

| VARIABLES | (1) | (2) | (3) | (4) | (5) |
|---|---|---|---|---|---|
| EA PGI (multi) | 0.133*** | | 1.135*** | | -7.871*** |
| | (0.003) | | (0.344) | | (1.598) |
| Parent edu. decile | | 0.069*** | -0.239 | | -0.122 |
| | | (0.002) | (0.212) | | (0.216) |
| Birth year | | | | 0.004*** | 0.004*** |
| | | | | (0.000) | (0.001) |
| EA PGI (multi) x Parent edu. decile | | | 0.006*** | | 1.082*** |
| | | | (0.002) | | (0.190) |
| EA PGI (multi) x Parent edu. decile x Birth year | | | | | -0.001*** |
| | | | | | (0.000) |
| Constant | -4.068*** | -7.539*** | -6.277*** | -6.607*** | -7.582*** |
| | (0.426) | (0.486) | (1.760) | (0.317) | (1.812) |
| Observations | 33,231 | 26,639 | 26,298 | 34,341 | 26,298 |
| R-squared | 0.101 | 0.085 | 0.134 | 0.027 | 0.135 |

b) Within family

| VARIABLES | (1) | (2) | (3) | (4) | (5) |
|---|---|---|---|---|---|
| Δ EA PGI (multi) | 0.082*** | | 0.074** | | 1.716 |
| | (0.007) | | (0.036) | | (4.393) |
| Parent edu. decile | | -0.001 | -0.002 | | -0.705 |
| | | (0.004) | (0.004) | | (0.473) |
| Birth year | | | | -0.000 | -0.003 |
| | | | | (0.000) | (0.002) |
| Δ EA PGI (multi) x Parent edu. decile | | | 0.001 | | -0.128 |
| | | | (0.004) | | (0.516) |
| Δ EA PGI (multi) x Parent edu. decile x Birth year | | | | | 0.000 |
| | | | | | (0.000) |
| Constant | -0.002 | 0.009 | 0.011 | 0.291 | 6.365 |
| | (0.007) | (0.033) | (0.033) | (0.621) | (4.036) |
| Observations | 6,343 | 5,824 | 5,684 | 6,609 | 5,684 |
| R-squared | 0.028 | 0.008 | 0.028 | 0.009 | 0.032 |

Note: Each between-family model includes controls for the first 20 principal components of the genetic data, and an interaction between birth year and sex. Each within-family model includes a control for sex difference within a twin pair. In the interaction models, all controls are interacted with the independent variables. Standard errors, shown in parentheses, allow for clustering at twin-pair level. *** p<0.01, ** p<0.05, * p<0.1

#### 6.2 Upper-secondary school performance

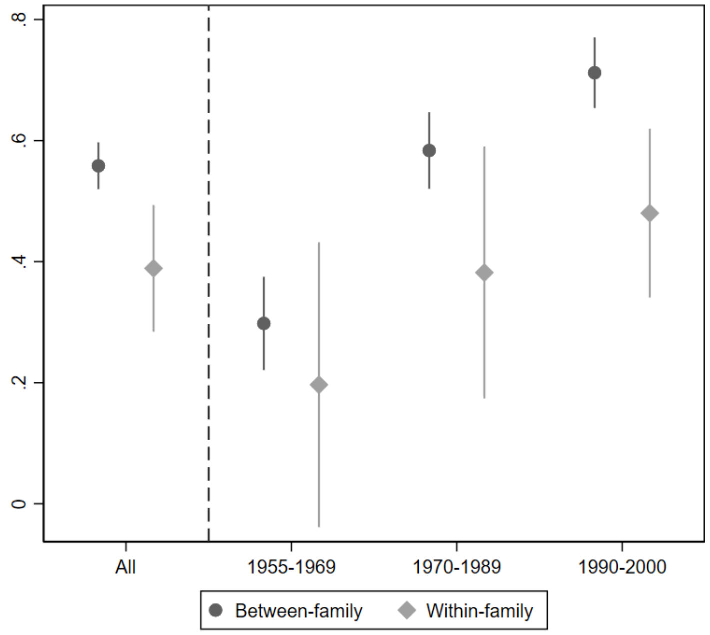

Figure A10: Effect of EA PGI on upper-secondary school performance by birth cohort

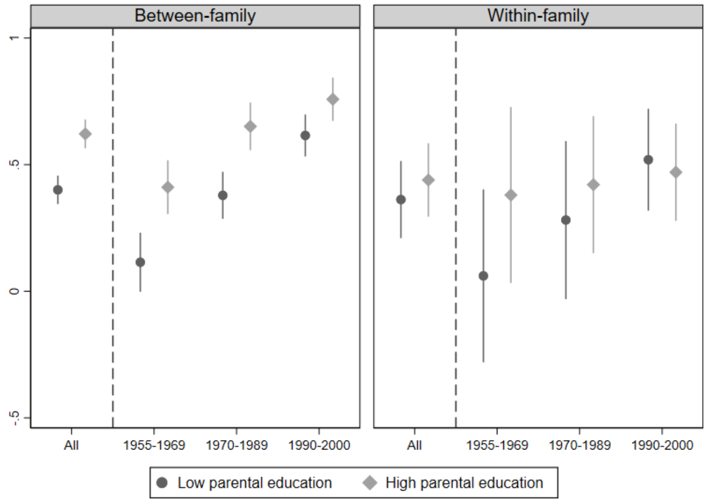

Figure A11: Effect of EA PGI on upper-secondary school performance by birth cohort, split on parental education

Table A16: Interaction between EA PGI and birth year for upper-secondary school performance, 1955–2000 (dummies)

a) Between family

| VARIABLES | (1) | (2) | (3) |
|---|---|---|---|
| EA PGI (multi) | 0.557*** | | 0.235*** |
| | (0.020) | | (0.045) |
| 1970-1989 | | 0.251*** | 0.363*** |
| | | (0.052) | (0.077) |
| 1990-2000 | | -0.298*** | -0.540*** |
| | | (0.051) | (0.076) |
| EA PGI (multi) x 1970-1989 | | | 0.285*** |
| | | | (0.051) |
| EA PGI (multi) x 1990-2000 | | | 0.420*** |
| | | | (0.050) |
| Constant | 7.080*** | 7.030*** | 7.093*** |
| | (0.060) | (0.045) | (0.060) |
| Observations | 24,599 | 25,583 | 24,599 |
| R-squared | 0.050 | 0.007 | 0.058 |

b) Within family

| VARIABLES | (1) | (2) | (3) |
|---|---|---|---|
| Δ EA PGI (multi) | 0.397*** | | 0.192* |
| | (0.052) | | (0.106) |
| 1970-1989 | | -0.286** | -0.320** |
| | | (0.138) | (0.139) |
| 1990-2000 | | -0.233** | -0.222* |
| | | (0.115) | (0.117) |
| Δ EA PGI (multi) x 1970-1989 | | | 0.170 |
| | | | (0.154) |
| Δ EA PGI (multi) x 1990-2000 | | | 0.299** |
| | | | (0.127) |
| Constant | -0.029 | 0.139 | 0.169* |
| | (0.047) | (0.098) | (0.099) |
| Observations | 4,559 | 4,812 | 4,559 |
| R-squared | 0.013 | 0.001 | 0.019 |

Note: Each between-family model includes controls for the first 20 principal components of the genetic data, and an interaction between birth year and sex. Each within-family model includes a control for sex difference within a twin pair. In the interaction models, all controls are interacted with the independent variables. Standard errors, shown in parentheses, allow for clustering at twin-pair level. *** p<0.01, ** p<0.05, * p<0.1

Table A17: Interaction between EA PGI and birth year for upper-secondary school performance, 1955–2000 (continuous)

a) Between family

| VARIABLES | (1) | (2) | (3) |
|---|---|---|---|
| EA PGI (multi) | 0.558*** | | -24.561*** |
| | (0.020) | | (2.798) |
| Birth year | | -0.010*** | -0.019*** |
| | | (0.001) | (0.002) |
| EA PGI (multi) x Birth year | | | 0.013*** |
| | | | (0.001) |
| Constant | 44.269*** | 26.963*** | 45.344*** |
| | (4.222) | (2.878) | (4.209) |
| Observations | 24,599 | 25,583 | 24,599 |
| R-squared | 0.045 | 0.003 | 0.053 |

b) Within family

| VARIABLES | (1) | (2) | (3) |
|---|---|---|---|
| Δ EA PGI (multi) | 0.397*** | | -17.873 |
| | (0.052) | | (13.525) |
| Birth year | | -0.005 | 0.016 |
| | | (0.003) | (0.014) |
| Δ EA PGI (multi) x Birth year | | | 0.009 |
| | | | (0.007) |
| Constant | -0.029 | 9.796 | -31.649 |
| | (0.047) | (6.239) | (27.577) |
| Observations | 4,559 | 4,812 | 2,049 |
| R-squared | 0.013 | 0.001 | 0.014 |

Note: Each between-family model includes controls for the first 20 principal components of the genetic data, and an interaction between birth year and sex. Each within-family model includes a control for sex difference within a twin pair. In the interaction models, all controls are interacted with the independent variables. Standard errors, shown in parentheses, allow for clustering at twin-pair level. *** p<0.01, ** p<0.05, * p<0.1

Table A18: Interaction between EA PGI, parental education, and birth year for upper-secondary school performance, 1955–2000 (dummies)

a) Between family

| VARIABLES | (1) | (2) | (3) | (4) | (5) |
|---|---|---|---|---|---|
| EA PGI (multi) | 0.965*** | | 0.388*** | | 0.401* |
| | (0.020) | | (0.100) | | (0.212) |
| Parent edu. decile | | 0.337*** | 0.170*** | | 0.170*** |
| | | (0.011) | (0.029) | | (0.029) |
| 1970-1989 | | | | 0.023 | -0.541** |
| | | | | (0.060) | (0.261) |
| 1990-2000 | | | | 0.105* | -0.244 |
| | | | | (0.057) | (0.251) |
| EA PGI (multi) x 1970-1989 | | | | | -0.136 |
| | | | | | (0.257) |
| EA PGI (multi) x 1990-2000 | | | | | 0.232 |
| | | | | | (0.241) |
| Parent edu. decile x 1970-1989 | | | | | 0.082*** |
| | | | | | (0.032) |
| Parent edu. decile x 1990-2000 | | | | | 0.068** |
| | | | | | (0.031) |
| EA PGI (multi) x Parent edu. decile | | | 0.047*** | | 0.045* |
| | | | (0.010) | | (0.025) |
| EA PGI (multi) x Parent edu. decile x 1970-1989 | | | | | 0.024 |
| | | | | | (0.030) |
| EA PGI (multi) x Parent edu. decile x 1990-2000 | | | | | -0.014 |
| | | | | | (0.029) |
| Constant | 11.186** | 2.199*** | 3.344*** | 4.878*** | 3.346*** |
| | (4.502) | (0.114) | (0.237) | (0.052) | (0.237) |
| Observations | 21,291 | 17,963 | 17,708 | 22,129 | 17,708 |
| R-squared | 0.139 | 0.081 | 0.168 | 0.025 | 0.168 |

b) Within family

| VARIABLES | (1) | (2) | (3) | (4) | (5) |
|---|---|---|---|---|---|
| Δ EA PGI (multi) | 0.397*** | | 0.054 | | 0.001 |
| | (0.052) | | (0.212) | | (0.502) |
| Parent edu. decile | | 0.030 | 0.033 | | -0.017 |
| | | (0.024) | (0.024) | | (0.061) |
| 1970-1989 | | | | -0.286** | -0.480 |
| | | | | (0.138) | (0.660) |
| 1990-2000 | | | | -0.233** | -0.724 |
| | | | | (0.115) | (0.573) |
| Δ EA PGI (multi) x 1970-1989 | | | | | -0.265 |
| | | | | | (0.680) |
| Δ EA PGI (multi) x 1990-2000 | | | | | 0.159 |
| | | | | | (0.571) |
| Parent edu. decile x 1970-1989 | | | | | 0.019 |
| | | | | | (0.079) |
| Parent edu. decile x 1990-2000 | | | | | 0.066 |
| | | | | | (0.069) |
| Δ EA PGI (multi) x Parent edu. decile | | | 0.044* | | 0.025 |
| | | | (0.026) | | (0.060) |
| Δ EA PGI (multi) x Parent edu. decile x 1970-1989 | | | | | 0.053 |
| | | | | | (0.082) |
| Δ EA PGI (multi) x Parent edu. decile x 1990-2000 | | | | | 0.019 |
| | | | | | (0.069) |
| Constant | -0.029 | -0.269 | -0.284 | 0.139 | 0.307 |
| | (0.047) | (0.191) | (0.193) | (0.098) | (0.520) |
| Observations | 4,559 | 4,639 | 4,505 | 4,812 | 4,505 |
| R-squared | 0.013 | 0.000 | 0.014 | 0.001 | 0.021 |

Note: Each between-family model includes controls for the first 20 principal components of the genetic data, and an interaction between birth year and sex. Each within-family model includes a control for sex difference within a twin pair. In the interaction models, all controls are interacted with the independent variables. Standard errors, shown in parentheses, allow for clustering at twin-pair level. *** p<0.01, ** p<0.05, * p<0.1

Table A19: Interaction between EA PGI, parental education, and birth year for upper-secondary school performance, 1955–2000 (continuous)

a) Between family

| VARIABLES | (1) | (2) | (3) | (4) | (5) |
|---|---|---|---|---|---|
| EA PGI (multi) | 0.558*** | | -24.904*** | | -32.127** |
| | (0.020) | | (2.936) | | (13.028) |
| Parent edu. decile | | 0.193*** | -10.498*** | | -10.426*** |
| | | (0.010) | (1.627) | | (1.619) |
| Birth year | | | | -0.010*** | -0.051*** |
| | | | | (0.001) | (0.007) |
| EA PGI (multi) x Parent edu. decile | | | 0.076*** | | 0.972 |
| | | | (0.010) | | (1.551) |
| EA PGI (multi) x Parent edu. decile x Birth year | | | | | -0.000 |
| | | | | | (0.001) |
| Constant | 44.269*** | 32.594*** | 108.353*** | 26.963*** | 107.396*** |
| | (4.222) | (4.312) | (13.388) | (2.878) | (13.326) |
| Observations | 24,599 | 20,812 | 20,513 | 25,583 | 20,513 |
| R-squared | 0.045 | 0.023 | 0.066 | 0.003 | 0.066 |

b) Within family

| VARIABLES | (1) | (2) | (3) | (4) | (5) |
|---|---|---|---|---|---|
| Δ EA PGI (multi) | 0.397*** | | 0.054 | | -12.112 |
| | (0.052) | | (0.212) | | (31.891) |
| Parent edu. decile | | 0.030 | 0.033 | | -4.682 |
| | | (0.024) | (0.024) | | (3.693) |
| Birth year | | | | -0.005 | -0.023 |
| | | | | (0.003) | (0.015) |
| Δ EA PGI (multi) x Parent edu. decile | | | 0.044* | | -0.784 |
| | | | (0.026) | | (3.851) |
| Δ EA PGI (multi) x Parent edu. decile x Birth year | | | | | 0.000 |
| | | | | | (0.002) |
| Constant | -0.029 | -0.269 | -0.284 | 9.796 | 44.920 |
| | (0.047) | (0.191) | (0.193) | (6.239) | (30.773) |
| Observations | 4,559 | 4,639 | 4,505 | 4,812 | 4,505 |
| R-squared | 0.013 | 0.000 | 0.014 | 0.001 | 0.019 |

Note: Each between-family model includes controls for the first 20 principal components of the genetic data, and an interaction between birth year and sex. Each within-family model includes a control for sex difference within a twin pair. In the interaction models, all controls are interacted with the independent variables. Standard errors, shown in parentheses, allow for clustering at twin-pair level. *** p<0.01, ** p<0.05, * p<0.1

#### 6.3 Income in early-middle adulthood

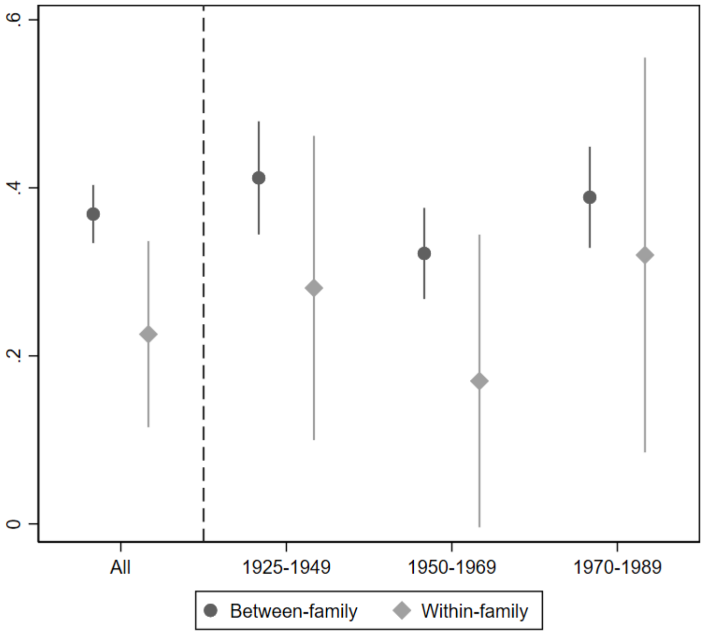

Figure A12: Effect of EA PGI on income by birth cohort

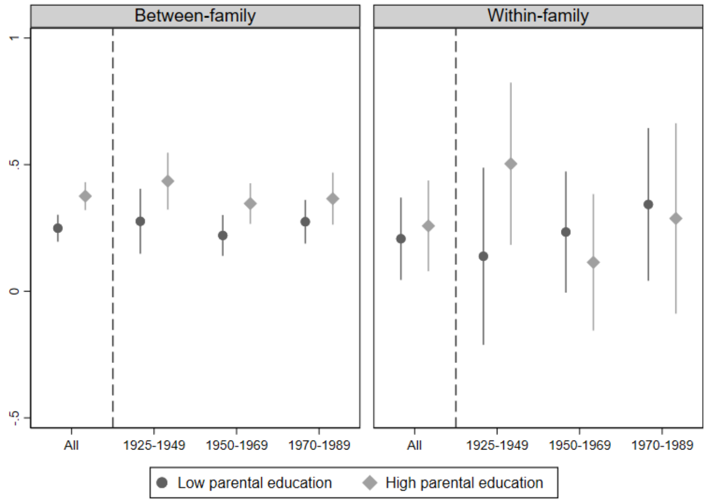

Figure A13: Effect of EA PGI on income by birth cohort, split on parental education

Table A20: Interaction between EA PGI and birth year for income in early-middle adulthood, 1940–1989 (dummies)

a) Between family

| VARIABLES | (1) | (2) | (3) |
|---|---|---|---|
| EA PGI (multi) | 0.369*** | | 0.401*** |
| | (0.018) | | (0.038) |
| 1950-1969 | | 0.340*** | 0.316*** |
| | | (0.045) | (0.064) |
| 1970-1989 | | 0.649*** | 0.441*** |
| | | (0.046) | (0.068) |
| EA PGI (multi) x 1950-1969 | | | -0.092** |
| | | | (0.044) |
| EA PGI (multi) x 1970-1989 | | | -0.013 |
| | | | (0.046) |
| Constant | 6.146*** | 6.078*** | 6.149*** |
| | (0.048) | (0.039) | (0.048) |
| Observations | 24,943 | 25,674 | 24,943 |
| R-squared | 0.033 | 0.011 | 0.038 |

b) Within family

| VARIABLES | (1) | (2) | (3) |
|---|---|---|---|
| Δ EA PGI (multi) | 0.222*** | | 0.247*** |
| | (0.056) | | (0.089) |
| 1950-1969 | | 0.097 | 0.085 |
| | | (0.115) | (0.117) |
| 1970-1989 | | 0.103 | 0.102 |
| | | (0.130) | (0.133) |
| Δ EA PGI (multi) x 1950-1969 | | | -0.073 |
| | | | (0.126) |
| Δ EA PGI (multi) x 1970-1989 | | | 0.022 |
| | | | (0.148) |
| Constant | -0.082 | -0.157* | -0.137* |
| | (0.051) | (0.082) | (0.083) |
| Observations | 4,264 | 4,408 | 4,264 |
| R-squared | 0.008 | 0.004 | 0.011 |

Note: Each between-family model includes controls for the first 20 principal components of the genetic data, and an interaction between birth year and sex. Each within-family model includes a control for sex difference within a twin pair. In the interaction models, all controls are interacted with the independent variables. Standard errors, shown in parentheses, allow for clustering at twin-pair level. *** p<0.01, ** p<0.05, * p<0.1

Table A21: Interaction between EA PGI and birth year for income in early-middle adulthood, 1940–1989 (continuous)

a) Between family

| VARIABLES | (1) | (2) | (3) |
|---|---|---|---|
| EA PGI (multi) | 0.369*** | | 0.979 |
| | (0.018) | | (2.372) |
| Birth year | | 0.017*** | 0.012*** |
| | | (0.001) | (0.002) |
| EA PGI (multi) x Birth year | | | -0.000 |
| | | | (0.001) |
| Constant | -16.730*** | -27.722*** | -16.567*** |
| | (3.532) | (2.342) | (3.532) |
| Observations | 24,943 | 25,674 | 24,943 |
| R-squared | 0.033 | 0.012 | 0.037 |

b) Within family

| VARIABLES | (1) | (2) | (3) |
|---|---|---|---|
| Δ EA PGI (multi) | 0.222*** | | 3.120 |
| | (0.056) | | (7.498) |
| Birth year | | -0.000 | -0.027** |
| | | (0.003) | (0.011) |
| Δ EA PGI (multi) x Birth year | | | -0.001 |
| | | | (0.004) |
| Constant | -0.082 | 0.157 | 53.242** |
| | (0.051) | (6.632) | (20.941) |
| Observations | 4,264 | 4,408 | 4,264 |
| R-squared | 0.008 | 0.004 | 0.012 |

Note: Each between-family model includes controls for the first 20 principal components of the genetic data, and an interaction between birth year and sex. Each within-family model includes a control for sex difference within a twin pair. In the interaction models, all controls are interacted with the independent variables. Standard errors, shown in parentheses, allow for clustering at twin-pair level. *** $p<0.01$, ** $p<0.05$, * $p<0.1$

Table A22: Interaction between EA PGI, parental education, and birth year for income in early-middle adulthood, 1940–1989 (dummies)

a) Between family

| VARIABLES | (1) | (2) | (3) | (4) | (5) |
|---|---|---|---|---|---|
| EA PGI (multi) | 0.369*** | | -0.042 | | -0.573 |
| | (0.018) | | (0.112) | | (0.379) |
| Parent edu. decile | | 0.191*** | 0.320*** | | 0.324*** |
| | | (0.011) | (0.044) | | (0.045) |
| 1950-1969 | | | | 0.340*** | 1.775*** |
| | | | | (0.045) | (0.413) |
| 1970-1989 | | | | 0.649*** | 2.127*** |
| | | | | (0.046) | (0.406) |
| EA PGI (multi) x 1950-1969 | | | | | 0.470 |
| | | | | | (0.408) |
| EA PGI (multi) x 1970-1989 | | | | | 0.591 |
| | | | | | (0.400) |
| Parent edu. decile x 1950-1969 | | | | | -0.160*** |
| | | | | | (0.047) |
| Parent edu. decile x 1970-1989 | | | | | -0.160*** |
| | | | | | (0.046) |
| EA PGI (multi) x Parent edu. decile | | | 0.045*** | | 0.105** |
| | | | (0.012) | | (0.043) |
| EA PGI (multi) x Parent edu. decile x 1950-1969 | | | | | -0.059 |
| | | | | | (0.047) |
| EA PGI (multi) x Parent edu. decile x 1970-1989 | | | | | -0.071 |
| | | | | | (0.046) |
| Constant | -16.730*** | 4.467*** | 3.311*** | 6.078*** | 3.271*** |
| | (3.532) | (0.113) | (0.390) | (0.039) | (0.393) |
| Observations | 24,943 | 19,970 | 19,750 | 25,674 | 19,750 |
| R-squared | 0.033 | 0.025 | 0.046 | 0.011 | 0.046 |

b) Within family

| VARIABLES | (1) | (2) | (3) | (4) | (5) |
|---|---|---|---|---|---|
| ∆ EA PGI (multi) | 0.222*** | | 0.357 | | -1.210 |
| | (0.056) | | (0.322) | | (1.093) |
| Parent edu. decile | | -0.042 | -0.041 | | -0.043 |
| | | (0.035) | (0.035) | | (0.108) |
| 1950-1969 | | | | 0.097 | -0.403 |
| | | | | (0.115) | (1.072) |
| 1970-1989 | | | | 0.103 | 0.332 |
| | | | | (0.130) | (1.052) |
| ∆ EA PGI (multi) x 1950-1969 | | | | | 1.623 |
| | | | | | (1.192) |
| ∆ EA PGI (multi) x 1970-1989 | | | | | 1.802 |
| | | | | | (1.195) |
| Parent edu. decile x 1950-1969 | | | | | 0.055 |
| | | | | | (0.122) |
| Parent edu. decile x 1970-1989 | | | | | -0.033 |
| | | | | | (0.120) |
| ∆ EA PGI (multi) x Parent edu. decile | | | -0.015 | | 0.169 |
| | | | (0.038) | | (0.124) |
| ∆ EA PGI (multi) x Parent edu. decile x 1950-1969 | | | | | -0.197 |
| | | | | | (0.136) |
| ∆ EA PGI (multi) x Parent edu. decile x 1970-1989 | | | | | -0.208 |
| | | | | | (0.137) |
| Constant | -0.082 | 0.274 | 0.272 | -0.157* | 0.240 |
| | (0.051) | (0.296) | (0.299) | (0.082) | (0.961) |
| Observations | 4,264 | 3,711 | 3,636 | 4,408 | 3,636 |
| R-squared | 0.008 | 0.003 | 0.008 | 0.004 | 0.010 |

Note: Each between-family model includes controls for the first 20 principal components of the genetic data, and an interaction between birth year and sex. Each within-family model includes a control for sex difference within a twin pair. In the interaction models, all controls are interacted with the independent variables. Standard errors, shown in parentheses, allow for clustering at twin-pair level. *** p<0.01, ** p<0.05, * p<0.1

Table A23: Interaction between EA PGI, parental education, and birth year for income in early-middle adulthood, 1940–1989 (continuous)

a) Between family

| VARIABLES | (1) | (2) | (3) | (4) | (5) |
|---|---|---|---|---|---|
| EA PGI (multi) | 0.369*** | | 2.439 | | -27.698* |
| | (0.018) | | (2.848) | | (14.871) |
| Parent edu. decile | | 0.197*** | 3.623** | | 4.074** |
| | | (0.011) | (1.721) | | (1.739) |
| Birth year | | | | 0.017*** | 0.037*** |
| | | | | (0.001) | (0.007) |
| EA PGI (multi) x Parent edu. decile | | | 0.040*** | | 3.614** |
| | | | (0.012) | | (1.742) |
| EA PGI (multi) x Parent edu. decile x Birth year | | | | | -0.002** |
| | | | | | (0.001) |
| Constant | -16.730*** | -30.068*** | -63.121*** | -27.722*** | -68.055*** |
| | (3.532) | (3.965) | (14.513) | (2.342) | (14.755) |
| Observations | 24,943 | 19,970 | 19,750 | 25,674 | 19,750 |
| R-squared | 0.033 | 0.026 | 0.046 | 0.012 | 0.046 |

b) Within family

| VARIABLES | (1) | (2) | (3) | (4) | (5) |
|---|---|---|---|---|---|
| Δ EA PGI (multi) | 0.222*** | | 0.357 | | -54.741 |
| | (0.056) | | (0.322) | | (45.074) |
| Parent edu. decile | | -0.042 | -0.041 | | 3.411 |
| | | (0.035) | (0.035) | | (4.825) |
| Birth year | | | | -0.000 | 0.013 |
| | | | | (0.003) | (0.021) |
| Δ EA PGI (multi) x Parent edu. decile | | | -0.015 | | 7.333 |
| | | | (0.038) | | (5.284) |
| Δ EA PGI (multi) x Parent edu. decile x Birth year | | | | | -0.004 |
| | | | | | (0.003) |
| Constant | -0.082 | 0.274 | 0.272 | 0.157 | -25.156 |
| | (0.051) | (0.296) | (0.299) | (6.632) | (41.274) |
| Observations | 4,264 | 3,711 | 3,636 | 4,408 | 3,636 |
| R-squared | 0.008 | 0.003 | 0.008 | 0.004 | 0.010 |

Note: Each between-family model includes controls for the first 20 principal components of the genetic data, and an interaction between birth year and sex. Each within-family model includes a control for sex difference within a twin pair. In the interaction models, all controls are interacted with the independent variables. Standard errors, shown in parentheses, allow for clustering at twin-pair level. *** p<0.01, ** p<0.05, * p<0.1

### 7 Miscellaneous

#### 7.1 Rolling regressions corresponding to main results

Figures A14 and A15 show rolling regressions, using 10-year windows. Figure A14 illustrates the interaction between the EA PGI and birth year, using the 1925–1989 sample. In other words, the main effect of the EA PGI is estimated for each consecutive birth year (± 5 years). Figure A15 plots the coefficient from the interaction between the EA PGI and parental education, using the 1940–1989 sample. Whereas the main results figure showed the interaction by splitting the sample by parental education, this graph shows the actual size of the interaction and how it changes over time.

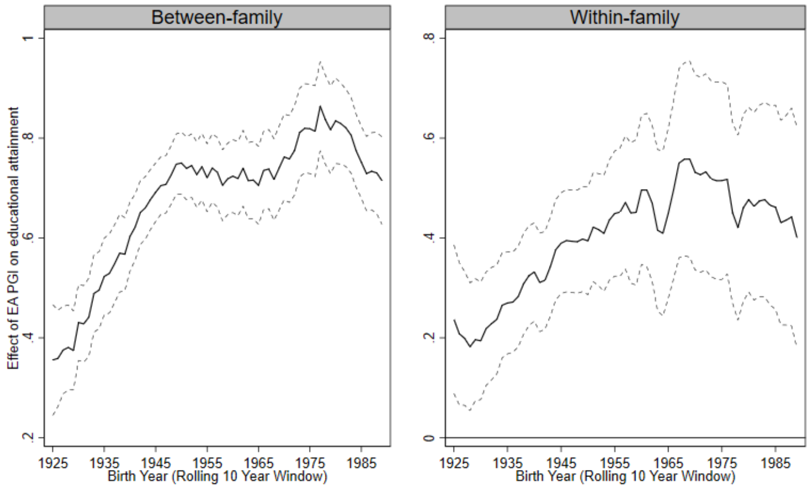

Figure A14: Relationship between EA PGI and educational attainment, 1925–1989 (rolling 10-year window)

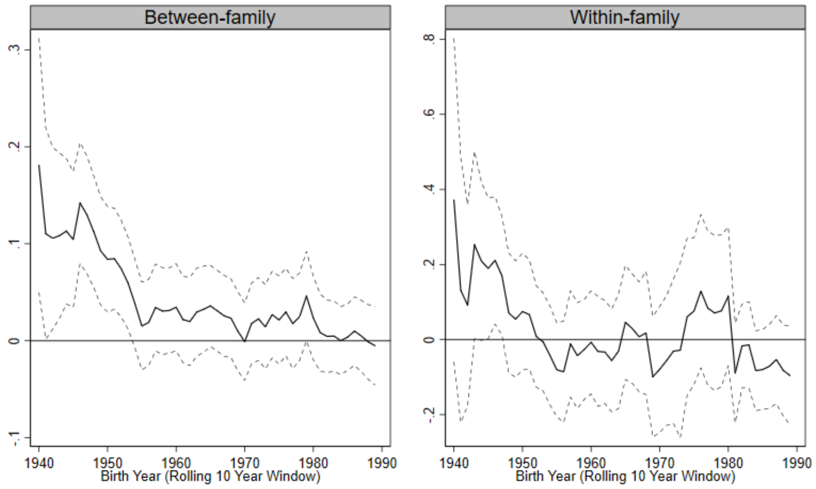

Figure A15: Interaction (size) between EA PGI and parental education, 1940–1989 (rolling 10-year window)
